# ATP-Powered Signaling between Synthetic and Living Cells

**DOI:** 10.1101/2025.08.13.670118

**Authors:** Soumya Sethi, Charu Sharma, Andreas Walther

**Author notes:** equal contribution (Soumya Sethi, Charu Sharma).

## Abstract

ATP is the energy currency of life and overabundant in the tumor microenvironment, where it has been suggested as a target for cancer therapy. We introduce ATP-dissipative delivery of DNA signals from synthetic cells to living cells by exploiting an ATP-driven reaction network that transiently ejects DNA Signal strands from the shielded synthetic cell interior to the extracellular medium of living cells. We customize the Signal for intracellular uptake or for extracellular instruction using a cytokine-ssDNA chimera that can trigger efficient intracellular downstream signaling programs. Our study discusses details of system design on a timer circuit and synthetic cell level, system integration challenges, and how ATP concentrations regulate the transient delivery. The strategy can be extended to deliver therapeutic oligonucleotides for applications in gene therapy and gene silencing. For cancer therapy, it can use naturally enhanced ATP levels to induce selective delivery of therapeutic oligonucleotides.

---

Interfacing synthetic cells (SC) and mammalian cells can facilitate development of programmable therapeutics that can sense, process, and respond to physiological changes^[1]^, create biohybrid systems for regenerative medicine^[2]^, enable targeted drug delivery[3], or allow synthetic tissue engineering.^[4]^ To establish effective communication between synthetic and living cells, direct cellular contact as well as (bio)chemical signaling systems have been considered.^[5]^ Contact-dependent communication relies on direct physical interactions at their interfaces, enabling specific signal exchange.^[6]^ These interactions can be engineered using programmable systems, relying on DNA self-assembly, which allow SCs to physically contact and recognize living cells.^[7]^ Besides this, the use of engineered signaling mechanisms^[8]^ has been demonstrated, for instance, SCs encapsulating transcription-translation machinery to instruct mammalian cells.^[9]^

ATP is a molecule of interest for communication among synthetic and mammalian cells due to its dual roles as both energy currency^[10]^ and signaling molecule.^[11]^ ATP is recognized as one of the main biochemical components of the tumor microenvironment (TME) with concentrations in the range of 50-200 µM.^[12–14]^ Depending on its concentration in the TME, the presence of ATP-hydrolyzing enzymes and receptors expressed by cancer and immune cells, tumor cell proliferation can be promoted or suppressed.^[15]^ With increased understanding of information exchange in TME, new therapeutic approaches can be developed to directly target extracellular ATP and the TME. In the context of SCs, this requires a biocompatible SC system that can release bioactive components of choice in response to ATP.

Building on ATP-fueled enzymatic reaction networks (ERNs)^[16–21]^, here we design an ATP-fueled DNA delivery system that transiently releases Signals (single-stranded ssDNA strand) into the extracellular media of cancer cells (Figure **1**). The system is composed of an ATP-driven ERN and SCs bearing the Signal to be delivered to target cells. The SC has the key function to shield the Signal against non-specific internalization by cells by harboring it in its interior. In the presence of ATP, the fuel-driven ERN is activated to temporally provide the Signal to the cells. Once ATP is consumed, the ATP-ERN resets the entire system to its initial state which involves recapturing of excess Signal by SCs. This enables controlled, ATP-dependent Signal delivery. We further extend this approach by coupling the Signal strand with a protein-DNA chimera for targeted delivery of cell-signaling cytokines. Importantly, this ATP-driven system actively consumes ATP, enabling self-regulated delivery responsive to ATP levels in the TME.

**Figure 1.**
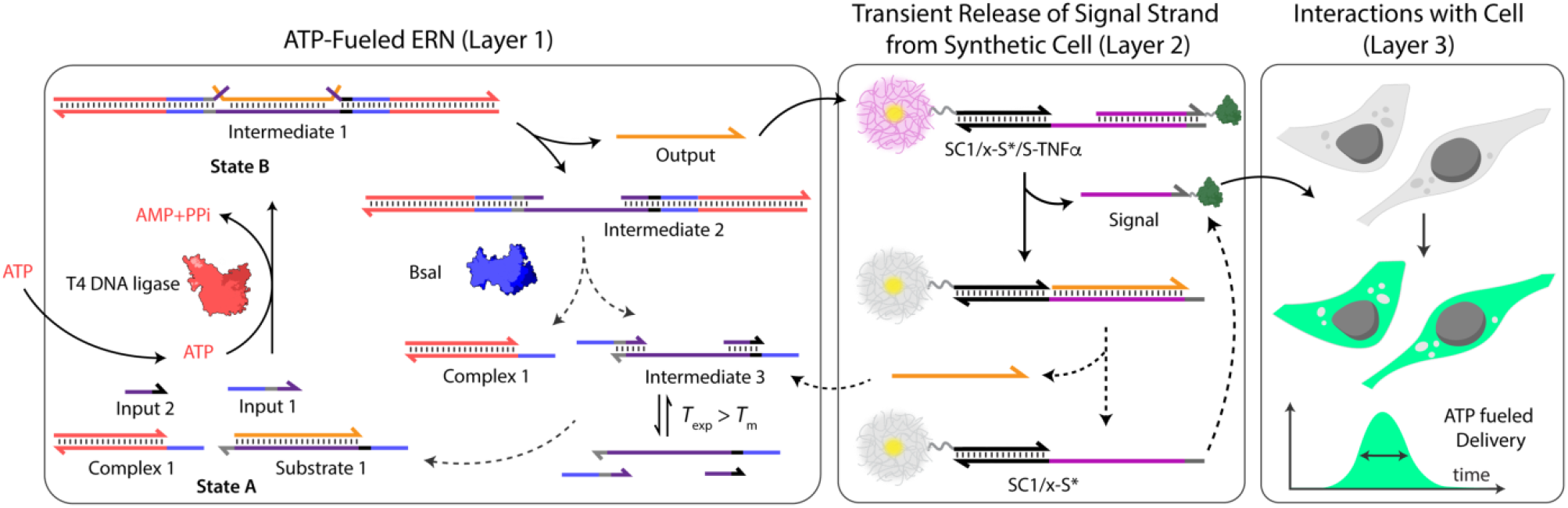
System design for controlled delivery of DNA in living cells via out-of-equilibrium extracellular medium.

In more detail, our system has three different layers: Layer 1 contains ATP-fueled ERN which operates as a molecular control program for transient release of Signal from SCs in Layer 2, which is followed by uptake of Signal by living cells in Layer 3. The ATP-driven ERN builds on our initial design^[16–21]^ of DNA-based ligation/restriction ERN that in the present configuration enables the ATP-powered transient release of a ssDNA that is recaptured once ATP is consumed in the ligation/restriction process. The clear separation with a shielding SC layer sets our design apart from recent work on ATP-instructed cell communication.^[22]^ In its initial deactivated State A, the ERN is composed of two double-stranded (ds) DNA complexes (Complex 1, Substrate 1) and two ssDNA (Input 1, Input 2; Figure **1**). In this state, Input 1 and Input 2 are incapable of displacing Output from Substrate 1 due to unstable hybridization of Input 1 (melting temperature (*T*_m_) = 32 ^°^C < experimental temperature (*T*_exp_) = 37 ^°^C, by NUPACK^[23]^) and Input 2 (*T*_m_ = 22 ^°^C < *T*_exp_ = 37 ^°^C) with longer strand of Substrate 1. However, addition of ATP powers covalent ligation of Substrate 1 with two molecules of Complex 1 and one molecule each of Input 1 and Input 2 with help of T4 DNA ligase, generating Intermediate 1. The system is pushed towards an activated transient state, where strand displacement from two sides now provides a strong thermodynamic push to release Output from Intermediate 1, finally generating Intermediate 2. The dual invasion strategy is essential for releasing an Output strand long enough to displace Signal in Layer 2.^[16,24]^ Concurrently, BsaI cleaves Intermediate 2 to regenerate Complex 1 and Intermediate 3. Because of low *T*_m_ (32 ^°^C < *T*_exp_ = 37 ^°^C), Intermediate 3 dissociates into Input 1 and Input 2, and reproduces Substrate 1 after re-hybridization with Output returning the system back to State A. Faster ligation than cleavage satisfies the kinetic boundary conditions required for the system to achieve a dynamic steady state (DySS) with an ATP-dependent lifetime of the Output before the restriction overtakes and brings the system back to its initial state. ^[16–21]^ Output is therefore generated in an uphill fashion and is transiently available in the system to perform downstream functions before its ultimately reassociated within Substrate 1.

We use the transiently released Output as a timer circuit to displace Signal from SC, hence conferring dissipative properties of the Output to the cell-instructive Signal. Once restriction overtakes ligation, Output prefers to regenerate Substrate 1 rather than residing with the SC (see NUPACK calculations in Figure **S1**). As a result, the SC can also recapture the Signal, which reforms the initial state of the system. The lifetime and amount of Signal released in the DySS depends not only on the ATP concentration but is also limited by the transient Output yield. This Signal can be designed to be internalized by mammalian cells, and is therefore lost for recovery, or it can be coupled to a cytokine to instruct cells extracellularly.

Interfacing this ATP-driven ERN with cells requires addressing several integration challenges. First, we require a cell-signaling strand that can conditionally be made available for cells and that remains “hidden” when not in the activated state to prevent unwanted signaling. We chose to immobilize it on SC based on a core-shell microgel^[25,26]^ that provides easy diffusive access for DNA strands for strand displacement, while at the same time shielding it in absence of Output messenger strand. Additionally, colloidal nature simplifies analytics in a CLSM during release and recapture. Second, for intracellular delivery, we require a Signal able to cross cell membranes in detectable amounts. Thus, we screened ssDNAs with different dyes, which are known to facilitate entry due to their hydrophobicity, and identified that Atto645N and Cy5-labeled ssDNA are readily taken up by HeLa cells and localize to the mitochondria, whereas Atto488-ssDNA is not taken up (Figure **S2**).^[27–30]^ Third, it is pivotal to suppress DNAse-mediated degradation of the DNA in ATP-driven ERN as well as in SCs by high nuclease concentrations present in typical cell culture medium containing fetal bovine serum (FBS).^[31]^ We optimized a dedicated cell culture medium to provide stability for over one day, whereas FBS-containing media shows significant degradation within 4 hours (Figure **S3**).

Before instructing HeLa cells, we turn to understanding coupling of ATP-ERN with SCs. For this, we functionalized micron-sized core-shell microgels^[25,26]^ as membrane-less SC chassis in their crosslinked PNIPAM-co-AA hydrogel shell with NH2-ssDNA through EDC-mediated coupling (84.2 wt% N-isopropylacrylamide (NIPAM); 10.4 wt% acrylic acid (AA); 1 wt% N,N’-methylenebis(acrylamide) (MBA), EDC = 1-Ethyl-3-(3-dimethylaminopropyl)carbodiimide; Figure **2**). This leads to a typical DNA grafting density of 3.3 × 10^6^ strands per SC or 132 ± 5 μmol strands per g of SC (Figure **S4**). Due to strong negative zeta potential of the SCs (ζ = −18 ± 2 mV) and the large size, such SCs cannot be internalized by cells within 24 h of incubation (Figure **S4**). Atto-647N-labeled Signal strand (S-Atto 647N) is immobilized with the help of a Linking strand (x-S*Q) to a final concentration of 3.75 μM (S-Atto 647N) on SCs with pendant x* ssDNA. CLSM confirms successful functionalization by appearance of bright fluorescence in Atto 647N channel (Figure **2b**). This strategy offers easy tuning of the amount of Signal on SCs.

**Figure 2.**
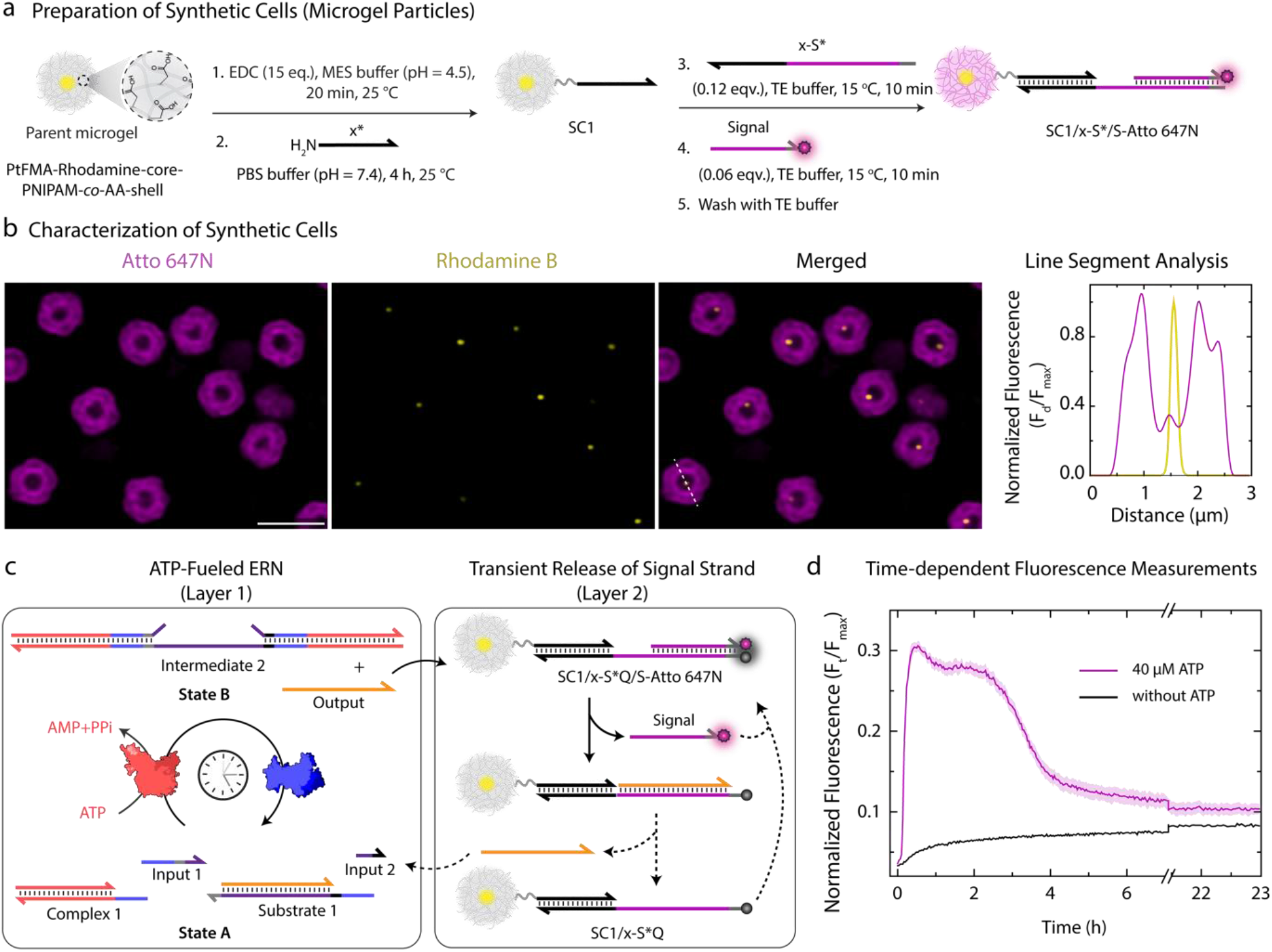
Preparation of DNA-functionalized SCs for transient release of Signal. (a) Scheme depicting stepwise preparation of SC1/x-S*/S-Atto 647N from parent SC. (b) CLSM characterization: SC1/x-S*/S-Atto 647N suspended in TE buffer (pH = 8.0) at 0.05 wt%. (c) Schematic representation of transient release of Signal from SC1/x-S*Q/S-Atto 647N via ATP-fueled ERN. (d) Time dependent FI changes demonstrate transient increase of fluorescence because of Signal strand release into medium upon ATP addition. Results are normalized with respect to 3.75 µM of S-Atto 647N (Signal) strands dissolved in DNA-Cell buffer which corresponds to the maximum fluorescence that can be observed in the system, average of two measurements. Shaded region depicts standard deviation (SD) of triplicates. Scale bar: 3 μm

Next, we coupled Signal-containing SCs to an ATP-fueled ERN to control the transient release of Signal in a dissipative fashion (Figure **2c**). Experimentally, ATP-fueled ERNs are optimized to operate at maximum efficiency in 1X Cut Smart buffer in Milli Q water. ^[17,24]^ However, to ensure the eventual coupling with cells, we simply prepared the 1X Cut Smart buffer in Cell buffer. We then checked the performance of this DNA-Cell buffer for transient Signal generation. To simplify monitoring via fluorescence intensity (FI) changes, we further exchanged the linking strand x-S* to a quencher-modified x-S*Q to monitor release of S-Atto 647N by appearance of fluorescence. Next, we assembled Signal-containing SCs (3.75 µM; 0.05 wt%) and ATP-driven ERN in DNA-Cell buffer at 37 ^°^C with 20 µM Complex 1, 5 µM Substrate 1, 10 µM Input 1 and Input 2 using 0.8 Weiss units (WU) µL^-1^ of T4 DNA ligase and controlled restriction takes over, and the Output, temporarily scavenged in SC on SC1/x-S*Q/Output, returns to Layer 1 and reforms Initial State A. This makes SC-x-S* available for Signal again causing its recapture dropping of FI. In previous works, we have shown how ATP can tune the lifetime of such systems and allow for reinitiation.^[16– 19]^ Figure **S5** shows exemplary tuning of lifetime by injecting 80 µM ATP, leading roughly to a doubling of the lifetime – as expected.

Understanding the amount of Signal transiently ejected from SCs is an important aspect. NUPACK simulations suggest that mixing 5 µM Output with 3.75 µM Signal complex present on the SCs (x*/x-S*/S = 7.5/7.5/3.75 µM) releases 3.5 µM Signal (Figure **S1, S6**), corresponding to 94% yield. Experimentally, direct addition of Output produces 3.52 µM Signal. In comparison, ATP-fueled Output generation, using 40 µM ATP, yields ca.1.47 µM (39 %) of Signal (details in Figure **S7**) due to the DySS nature of the system. From previous experiments for cellular uptake of dye-labelled ssDNA (Figure **S2**), we know that this concentration is sufficient to efficiently transfer Signal into HeLa cells within 2.5 h. Note that a minimal increase of ca. 6% in the absence of ATP is caused by unavoidable leakage between Substrate 1 and the Signal complex in the SCs (Figure **S7**). However, this increase in fluorescence is slow and insufficient to cause any observable cellular internalization.

Building on the understanding of the transient ATP-driven release from SCs, we next combined it with cellular delivery using HeLa cells as model cell line (Figure **3**). CLSM helps to understand details of the transfer process (Figure **3b**). We prepared Signal-containing SCs, but omitted the quencher on the Linker strand, enabling tracking of Signal transfer from SCs to the cells all the way through the medium upon ATP addition. With SCs in hand, we integrated the same ATP-fueled ERN as above and triggered it with 40 µM ATP. Before adding ATP, SCs show high fluorescence due to the bonded Signal, and no fluorescence can be detected within cells marked with white dotted lines (Figure **3b-c**). However, within 18 minutes of ATP addition, FI on SCs decreases by 90%, and a concurrent FI increase can be detected in the medium, attaining a transient maximum FI (Figure **3c**). Notably, during the next 2.5 h, FI in the medium decreases, whereas FI of the cells increases, and FI of SCs remains stable. This clearly demonstrates that in this DySS, Signal is being transferred of the Output from the ATP fueled process into cells. Once ATP is consumed (ca. at 2.5 h, Figure **3d**), SCs begin to recover their FI by recapturing excess Signal from the medium. A partial recovery of only 37% in FI on the SCs confirms partial internalization of Signal by cells, which reaches a similar FI at the end. Note that S-Atto 647N localizes to the mitochondria of cells. Upon comparing FI decrease on SCs with FI increase of the medium at 24 h of ATP addition, we estimate the amount of Signal internalized within cells during this time to be 2.06 µM or 1.24 × 10^11^ Signal strands per cell (see Figure **S7** for further details). A control system without ATP shows only minor Signal transfer due to minor strand displacement leakage (Figure **3c** (left), Figure **S8**). Cell viability remains high with an overall decrease of only 12% after 24 h (Figure **S11**).

**Figure 3.**
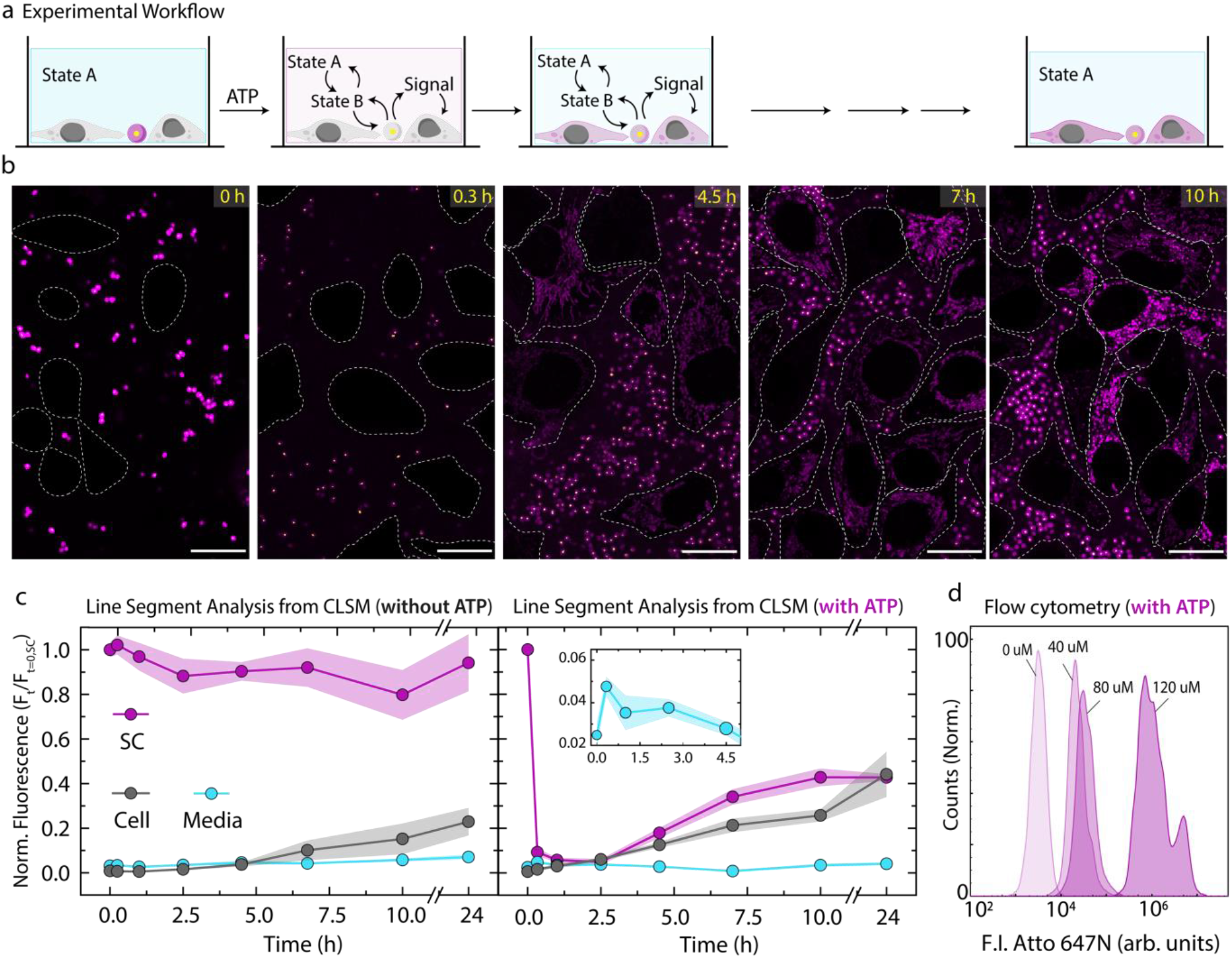
ATP-fueled signal transfer from SCs to HeLa cells. (a) Scheme of experimental workflow where Layer1 and Layer2 are installed within extracellular medium of cells marked by initial State A. State B is acquired upon ATP addition which marks transient release of Signal strands from SCs into medium to be uptaken by cells. (b) CLSM imaging of ATP-fueled delivery of Signal from SCs into cells. Dotted lines define the cell boundary. (c) Time-dependent FI changes on SCs, in medium and within cells without and with ATP. Inset shows zoomed in region of FI changes in media. The fluorescence values represent the maximum intensity acquired from line segment analysis and normalized with respect to maximum fluorescence observed on SCs. Results represent average contribution from 5 different regions from 3 replicates, shaded regions depict SD. (d) Flow cytometer measurement of cells in presence of ATP fueled ERN networks at various ATP concentrations to study release and uptake of dye-labelled Signal. Scale bars: 20 μm.

More importantly, we quantitatively assessed the percentage of cells that incorporate the Signal under varying ATP concentrations, from 0 to 120 µM, using flow cytometry at 30 min time point. In absence of ATP, cells exhibit minimal uptake of Signal, resulting in lowest Atto 647N FI. As expected, at ATP concentrations of 40, 80 and 120 µM, the percentage of positive cells is high with a slight increase from 97.6% to 99.9% (Figure **S10**). However, the extent of delivered Signal is strongly different as the FI per cell correspondingly increases with higher ATP concentrations. (Figure **4d**). Furthermore, we analyzed cells at fixed ATP concentrations (40 µM, 80 µM) across different time points (0.5 h, 2 h, 4 h). As time progresses, cells take up more Signal, and the FI within cells increases accordingly (Figure **S11, S12**).

**Figure 4.**
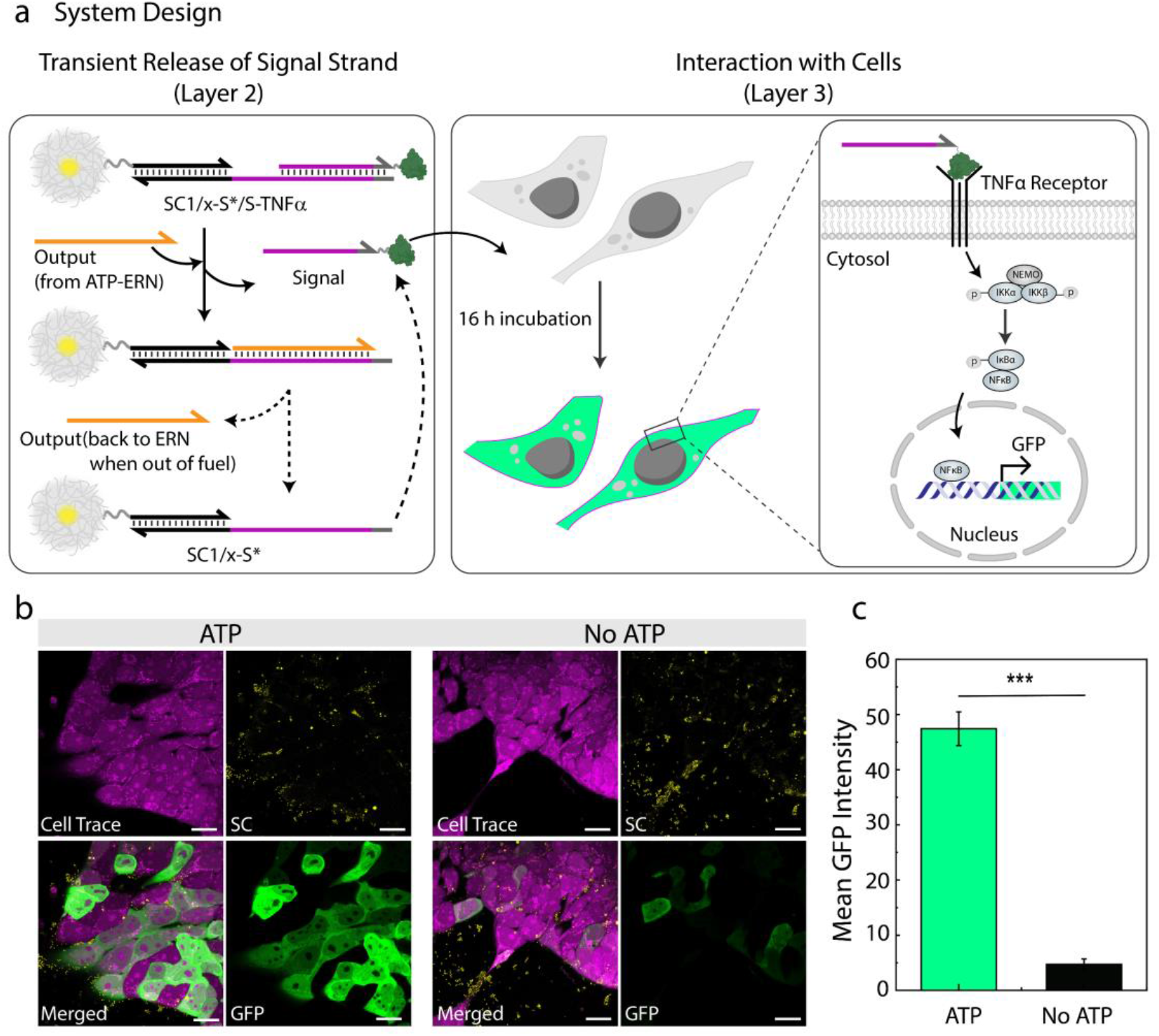
Cellular response to ATP-fueled release of TNFa-linked ssDNA Signal. (a) Simplified scheme illustrating the incubation of cells and SCs in ATP-fueled environments. (b) CLSM imaging of GFP fluorescence expressed by the TNFa-RCL in absence and presence (40 µM) of ATP after 16 h. (c) Quantified mean GFP intensity from CLSM images of cells in the presence and absence of ATP. The results are presented as the mean ± SEM, n= 72 cells from 3 replicates. Statistics: Unpaired, two-tailed student’s t-test. Scale bars: 20 μm

Directing cellular behavior with the ATP-fueled ERN system is critical for applications in cancer biology^[32]^, where modulation of extracellular signals provides an entry point for therapy. Thus, we investigated whether our system can be used to trigger programmed release of a cytokine to affect cellular function. To do so, we modified our approach for Signal strand and conjugated TNFα (tumor necrosis factor) protein to it. TNFα is an important TME signal to promote cancer cell death and neoantigen presentation, overall promoting immune response in cancer. To understand whether ATP-dissipating signaling can be exploited for extracellular signaling, we employed a reporter cell line, i.e., NF-κB/293/GFP-Luc Transcriptional Reporter Cell Line (TNFα-RCL) to analyze nuclear factor Kappa B (NF-κB) pathway. In the presence of TNFα ligand, TNFα-RCL generates GFP in a concentration dependent fashion, readily detected by CLSM (Figure **4a**).^[33]^ We first examined the effect of native TNFα on TNFα-RCL and observe that TNFα-RCL produce GFP fluorescence (Figure **S13**). Then, we accessed the efficacy of TNFα-ssDNA conjugate by incubating with TNFα-RCL in DNA-Cell Buffer. Indeed, TNFα-RCL produces GFP fluorescence (Figure **S14**), confirming that TNFα-RCL thrives in DNA-Cell Buffer and TNFα-ssDNA conjugate signals similarly as native TNFα. Next, we investigated the time point of GFP production in TNFα-RCL, noting that 16 h after dosing is sufficient for GFP production (Figure **S15**). Finally, we assembled our SC systems with S-TNFα together with upstream ATP-fueled ERN along with TNFα-RCL (Figure **4a**). Adding ATP as a fuel leads to a transient S-TNFα release, indeed triggering the NF-κB pathway. This results in a 10-fold higher GFP fluorescence with respect to the control (no ATP) where GFP expression is close to zero (Figure **4b-c**). These findings demonstrate that our ATP-driven reaction network is both robust and adaptable, capable of functioning in extracellular environment of cells and can be engineered to release protein-linked DNA strands on demand to trigger cellular pathways.

In summary, we have demonstrated an ATP-powered SC system to control transient delivery of DNA signals from shielded SC configuration to living cells. ATP consumption drives Signal release and its eventual recapture, imparting temporal control and dissipative behavior to the signaling process. Dissipative behavior returns the system back to its original state once fuel is exhausted, giving rise to homeostatic control mechanisms. For successful operation of longer ERNs, we customized media to maintain stability of all DNA-based components. Since the rate of DNA uptake by cells is a cell-intrinsic parameter, the rate of DNA release and recapture from upstream ERN needs to be programmed to match uptake kinetics. This signal can either be engineered for uptake by mammalian cells – rendering it unrecoverable – or linked to a cytokine to act externally and guide cellular behavior, where the reporter cell line provided clear evidence for successful cytokine instruction. Due to modularity of our design, therapeutic oligos such as siRNA, miRNA and antisense oligos may be delivered into cells to enable advanced applications in gene therapy and gene silencing. Fuel concentrations of 50-200 µM have been detected in TME, which is commensurate with ATP-ERN system presented here. A future realistic SC-based delivery platform needs to immobilize enzymes and all other components of the ERN inside a fully self-contained SC to allow for instance peritumoral application.

## Supporting Information

The authors have cited additional references within the Supporting Information. ^[26,33–35]^

## Acknowledgements

This project has received funding from the European Research Council (ERC) under the European Union’s Horizon 2020 research and innovation program (M3ALI: 101001638), as well as from the CoM2Life start-up funds of JGU. AW acknowledges funding via a Gutenberg Research Professorship underpinning his Life-Like Materials Program.

## References

[1] A. Joesaar, S. Yang, B. Bögels, A. Van Der Linden, P. Pieters, B. V. V. S. P. Kumar, N. Dalchau, A. Phillips, S. Mann, T. F. A. De Greef, Nat. Nanotechnol. 2019, 14, 369–378.

[2] Y. Sümbelli, A. F. Mason, J. C. M. Van Hest, Adv. Biol. 2023, 7, 2300149.

[3] J. A. M. Steele, J.-P. Hallé, D. Poncelet, R. J. Neufeld, Adv. Drug Deliv. Rev. 2014, 67–68, 74–83.

[4] M. H. M. E. Van Stevendaal, J. C. M. Van Hest, A. F. Mason, ChemSystemsChem 2021, 3, e2100009.

[5] X. Wang, L. Tian, H. Du, M. Li, W. Mu, B. W. Drinkwater, X. Han, S. Mann, Chem. Sci. 2019, 10, 9446–9453.

[6] L. Li, S. Liu, C. Zhu, S. Shao, F. Yang, Q. Liu, W. Tan, Angew. Chem. Int. Ed. 2025, 64, e202503903.

[7] M. Wang, H. Nan, M. Wang, S. Yang, L. Liu, H.-H. Wang, Z. Nie, Nat. Commun. 2025, 16, 2410.

[8] V. Mukwaya, S. Mann, H. Dou, Commun. Chem. 2021, 4, 161.

[9] Ö. D. Toparlak, J. Zasso, S. Bridi, M. D. Serra, P. Macchi, L. Conti, M.-L. Baudet, S. S. Mansy, Sci. Adv. 2020, 6, eabb4920.

[10] S. Hwang, M. Kim, A. P. Liu, ChemPlusChem 2024, 89, e202400138.

[11] I. Novak, Physiology 2003, 18, 12–17.

[12] S. Gilbert, C. Oliphant, S. Hassan, A. Peille, P. Bronsert, S. Falzoni, F. Di Virgilio, S. McNulty, R. Lara, Oncogene 2019, 38, 194–208.

[13] F. Di Virgilio, E. Adinolfi, Oncogene 2017, 36, 293–303.

[14] P. Pellegatti, L. Raffaghello, G. Bianchi, F. Piccardi, V. Pistoia, F. Di Virgilio, PLoS ONE 2008, 3, e2599.

[15] G. Burnstock, F. Di Virgilio, Purinergic Signal. 2013, 9, 491–540.

[16] J. Deng, A. Walther, Nat. Commun. 2021, 12, 5132.

[17] J. Deng, A. Walther, J. Am. Chem. Soc. 2020, 142, 685–689.

[18] J. Deng, A. Walther, Chem 2020, 6, 3329–3343.

[19] C. Sharma, A. Sarkar, A. Walther, Chem. Sci. 2023, 14, 12299–12307.

[20] Y. Xu, Y. Luo, X. Lu, J. Ye, Z. Chen, Y. Hu, C. Shen, B. Zhao, E. Kou, J. Deng, C. Fan, H. Zhang, H. Zhang, J. Am. Chem. Soc. 2025, DOI 10.1021/jacs.5c08925.

[21] J. N. Zadeh, C. D. Steenberg, J. S. Bois, B. R. Wolfe, M. B. Pierce, A. R. Khan, R. M. Dirks, N. A. Pierce, J. Comput. Chem. 2011, 32, 170–173.

[22] J. Deng, A. Walther, J. Am. Chem. Soc. 2020, 142, 21102–21109.

[23] K. Han, D. Go, T. Tigges, K. Rahimi, A. J. C. Kuehne, A. Walther, Angew. Chem. Int. Ed. 2017, 56, 2176–2182.

[24] K. Han, D. Go, D. Hoenders, A. J. C. Kuehne, A. Walther, ACS Macro Lett. 2017, 6, 310–314.

[25] A. S. Walsh, H. Yin, C. M. Erben, M. J. A. Wood, A. J. Turberfield, ACS Nano 2011, 5, 5427–5432.

[26] J. Mikkilä, A.-P. Eskelinen, E.H. Niemelä, V. Linko, M. J. Frilander, P. Törmä, M. A. Kostiainen, Nano Lett. 2014, 14, 2196–2200.

[27] V. J. Schüller, S. Heidegger, N. Sandholzer, P. C. Nickels, N. A. Suhartha, S. Endres, C. Bourquin, T. Liedl, ACS Nano 2011, 5, 9696–9702.

[28] A. Lacroix, E. Vengut-Climent, D. De Rochambeau, H. F. Sleiman, ACS Cent. Sci. 2019, 5, 882–891.

[29] J. Fern, R. Schulman, ACS Synth. Biol. 2017, 6, 1774–1783.

[30] G. Liu, L. Yang, G. Chen, F. Xu, F. Yang, H. Yu, L. Li, X. Dong, J. Han, C. Cao, J. Qi, J. Su, X. Xu, X. Li, B. Li, Front. Pharmacol. 2021, 12, 735446.

[31] N. Ramani, C. A. Figg, A. J. Anderson, P. H. Winegar, E. Oh, S. B. Ebrahimi, D. Samanta, C. A. Mirkin, Adv. Mater. 2023, 35, 2301086.

