## Supplementary material for "ATP-Powered Signaling between Synthetic and Living Cells": Sup Info

#equal contribution

[b] present address:
Delft University of Technology
Kavli Institute of Nanoscience
Department of Bionanoscience

### Experimental Materials and Methods

#### 1. Materials

All chemicals and reagents were purchased from Sigma Aldrich or Merck and used without further purification unless otherwise stated:

2,2,2-trifluoroethyl methacrylate (ABCR GMBH, 99 %), N-isopropylacrylamide (97 %), 2,2'-azobis(2-methylpropionamidine) dihydrochloride (ABCR GMBH, 96 %), N,N'-methylenebis(acrylamide) (99%), divinylbenzene (DVB, 80 %), acryloxyethyl thiocarbamoyl Rhodamine B (POLYSCIENCES, INC.), Al2O3 90 neutral (CARL ROTH GMBH), 1-Ethyl-3-(3-dimethylaminopropyl)carbodiimide (ABCR GMBH, 98 %), potassium persulfate (≥99 %), acrylic acid (Acros Organics, 99.5 %), disodium ethylenediaminetetraacetate dehydrate (biology grade), tris(hydroxymethyl)aminomethane hydrochloride (Trizma buffer substance pH=8), 2-(N-morpholino) ethanesulfonic acid, sodium chloride (99 %), magnesium chloride (99 %), phosphate buffered saline (PBS, pH = 7.4), Hoechst 34580 (Thermo Fisher, 98%), CellMask™ green plasma membrane dye (Thermo Fisher, 98%), Human TNF-alpha solution (Genscript, Z01001), NAP-10 desalting column (Cytiva, 17-0854-02), CellTrace™ violet (Thermo Fisher)

T4 DNA Ligase Storage Buffer (Promega): 10 mM Tris-HCl (pH 7.4 at 25 °C), 50 mM KCl, 1 mM dithiothreitol (DTT), 0.1 mM EDTA, 50 % glycerol.

BsaI-HF® v2 storage buffer (NEB): 10 mM Tris-HCl, 200 mM NaCl, 1 mM DTT, 0.1 mM EDTA, 200 µg/mL BSA, 50% glycerol.

1X NEB CutSmart® Buffer: 50 mM potassium acetate, 20 mM Tris-acetate, 10 mM magnesium acetate, 100 µg mL-1 BSA. Annealing Buffer: 10 mM Tris-HCl (pH 8.0), 50 mM NaCl, 10 mM MgCl2.

Cell buffer: 1X Insulin-Transferrin-Selenium (ITS-G, 100X, Gibco™), 1% MEM NEAA (Non Essential Amino Acid Solution, 100X, without L-Glutamine, PAN Biotech), 1% ROTI®Cell HEPES solution (1 M, Carl Roth), 1% Penicillin-Streptomycin (5000 U/mL, Gibco™) in MEM Eagle with: EBSS (Earle's Balanced Salt Solution), stable glutamine, 2.2 g/L NaHCO3 (PAN Biotech).

DNA-Cell buffer: 1X NEB CutSmart® Buffer in Cell buffer.

MilliQ water was used throughout all experiments unless otherwise stated.

All oligonucleotides (except amine-modified oligonucleotides) were purchased from Integrated DNA Technologies Inc. (IDT) and Biomers GmbH (as listed below in Table S1). The oligonucleotides received were dissolved in 1X TE buffer, pH-= 8.0 (Thermo Fischer). Amine-modified oligonucleotides were synthesized following a well-defined procedure (as described in section 3.4).

#### 2. General Characterization Methods and Instruments

##### *2.1 DLS measurements*

DLS measurements were performed on the LS Instruments NanoLab 3D at 25 °C operating with a red laser (λ = 685 nm) and a scattering angle of Θ = 90 ° using standard disposable PS cuvettes (BRAND GmbH & Co. KG). The distributions of the hydrodynamic radii were obtained by a CONTIN mode analysis

##### *2.2 Zeta-Potential*

(ζ-potential) of microgels was measured using a Zetasizer Nano ZS (Malvern Panalytical) at 25 °C. All samples were measured in triplicate in disposable folded capillary cells.

##### *2.3 Statistical Analysis*

Each experiment was performed at least three independent times. Statistical results were analyzed using Origin 2023b and Prism 8.0. Comparisons between two groups were conducted using unpaired two-tailed Student's t test (Figure 4c). For all analyses, *** indicates $P < 0.001$ and was considered significant.

##### *2.4 Fluorescence and Brightfield Microscopy*

Fluorescence and Brightfield Microscopy was performed on EVOS™ M7000 Imaging System equipped with DAPI, GFP, RFP and Cy5 filter cubes using 4×, 20×, and 40× objectives.

##### *2.5 Confocal Laser Scanning Microscopy*

Confocal laser scanning microscopy (CLSM) was performed on Leica Stellaris 5 microscope (LasX v4.3.0.24308) with four laser lines and three HyD S detectors using plan-apochromat objectives (63×, 1.40 numerical aperture, oil immersion).

##### *2.6 DNA Concentrations*

DNA concentrations were determined using a DeNovix-S-06873 (DeNovix OS 0.8.1 v4.1.5) spectrophotometer with a standard value of 33 µg/OD260.

##### *2.7 Temperature Controlled fluorescence measurements*

The temperature-controlled fluorescence measurements were performed on a TECAN (SPARK control v3.1) microplate plate reader using Corning® 384-Well black polystyrene plate with non-binding surface. Excitation and emission wavelengths for Atto 488 are 485 nm and 535 nm and for Atto 647N are 620 nm and 679 nm respectively.

##### *2.8 Flow Cytometry*

Flow Cytometry experiments were performed on a Novocyte Quanteon (Agilent, NovoExpress v.1.6.0) with 4 excitation lasers (violet 405 nm, blue 488 nm, yellow-green 561 nm and red 640 nm) and 16 fluorescence detectors. Flow Cytometry data was analyzed using FLOWJO (v10.9) software.

#### 3. Methods

##### *3.1 Synthesis of surfactant-free, poly(2,2,2-trifluoroethyl methacrylate)-Rhodamine labeled core particles*

The synthesis is analogous to a previous report.[1,2] Divinylbenzene (DVB) and 2,2,2-trifluoroethyl methacrylate (tFMA) were purified using column chromatography ($Al_2O_3$, neutral). The initiator potassium persulfate (KPS, 159.49 mg, 590 µmol) was dissolved in deionized water (45 mL), degassed by bubbling with nitrogen gas (25 min) and thermostated at 70 °C for 15 min. To a solution of DVB (46.2 mg, 354.9 µmol) and tFMA (1.98 g, 11.8 mmol), a solution of acryloxyethyl thiocarbamoyl Rhodamine B (1 mg) and N-isopropylacrylamide (NIPAM, 150 mg, 1.3 mmol) was added in water (4.5 mL). The mixture was ultrasonicated for 2 minutes and degassed by bubbling with nitrogen gas (10 min). The resulting mixture was added dropwise (over a period of 5 min) to the initiator solution starting the polymerization. The reaction mixture was stirred at 70 °C for 6 h (stirring rate = 600 rpm). The resulting dispersion was filtered while hot and dialyzed against deionized water (MWCO 8000 Da; solid content = 35.6 mg/mL by freeze-drying).

##### *3.2 Synthesis of the PNIPAM shell to yield PtFMA-Rhodamine labelled-core-PNIPAM-co-GMA-shell microgel (MGs) particles.*

The reaction conditions was adapted from a previous report[3]: NIPAM (3.37 g, 29.8 mmol, 84.2 wt %) was dissolved in deionized water (100 mL) together with co-monomer acrylic acid (AA, 418 mg, 398 µL, 5.8 mmol, 10.4 wt%), and the cross-linker N,N'-methylenebis(acrylamide) (MBA, 41.8 mg, 271 µmol, 1 wt %). The core particles (4.2 mL, 77 mg/mL, 170 mg solid content, 4.25 wt %) were added and the mixture was degassed for 30 min and heated to 45 °C. The initiator KPS (76.2 mg, 0.28 mmol) was dissolved in water (30 mL) and degassed (15 min). The polymerization was initiated by dropwise addition of initiator solution to a heated reaction mixture and stirred (stirring rate = 450 rpm). Immediately following initiation, a temperature ramp from 45 to 65 °C was applied to the solution at an approximate ramp rate of 20 °C/h. The stirring rate was increased to 600 rpm and reaction was allowed to run for 4 hours. The resulting core/shell MG particles (amount of AA moieties assuming full conversion = 1450 µmol per g of MG) were filtered while hot and purified by dialysis (MWCO 8000 Da) against deionized water. The resulting core-shell MGs were further purified via centrifugation (5 × 25 min, 11000 rpm, 15 °C, replacement of the supernatant with Milli Q water per centrifugation step).

##### *3.3 Functionalization of MG with amine-modified DNA.*

The MG suspension was redispersed in 2-(N-morpholino)ethanesulfonic acid (MES) buffer (10 mM, pH = 4.5) via centrifugation (20 min, 9000 rpm, 15 °C, replacement of the supernatant with MES buffer). A two-step reaction achieves the functionalization of DNA on MGs. In the first step, activation of carboxyl acid groups in MG shell (120 µL, 1.943 mg/mL of MGs, 0.34 µmol of COOH groups) was carried out by stirring with EDC (1.1 mg, 7.2 µmol, 25 equiv. with respect to COOH groups) dissolved in 40 µL of MES buffer (10 mM MES, pH = 4.5) for 12 minutes at 25 °C. Finally, amine-modified DNA (0.08 µmol, 0.23 equiv.) dispersed in 350 µL PBS buffer (pH = 7.4) was added to activated MGs prepared in first step and the mixture was stirred for 4.5 h at 25 °C. The DNA functionalized MGs were purified via centrifugation (2 × 3 min, 8000 rpm, 25 °C, replacement of the supernatant with TE buffer (pH = 8.0), per centrifugation step).

##### *3.4 Synthesis and purification of amine-modified oligonucleotide sequences.*

The oligonucleotides were synthesized at 10 µmol scale employing the standard solid phase β-cyanoethyl-phosphoramidite chemistry in trityl-on mode. The DNA phosphoramidites (DMT (dimethoxytrityl)-dT, DMT-dA(bz), DMT-dG(dmf) and DMT-dC(ac)) were diluted to 50 mM with dry acetonitrile and synthesis occurred from the 3' towards the 5' end of the oligonucleotides on packed solid phase columns.

Cleavage of the oligonucleotides (DMT-on) from the solid support and base deprotection was achieved in one step to ensure optimal yields. The 34 µmol/g controlled pore glass (CPG) solid support was treated with 10 mL of ice-cold ammonia solution (30-32 % $NH_3$) overnight at room temperature to detach the DNA from the CPG support. The cleaved DNA (DMT-on) in ammonia was diluted with 10 mL of disodium phosphate buffer (75 mM containing 1 mM EDTA, pH = 8.3) and the crude product was obtained upon freeze drying. The obtained DNA was redispersed in MilliQ water and purified by preparative reverse phase-HPLC (RP-HPLC) followed by freeze drying to remove the solvent.

The DMT group was cleaved from the purified product by making a 2 wt % solution of the dry DNA in NaOAc/HOAc buffer (200 mM, pH 4.0, 200 mM NaCl) and heating the mixture to 50 °C for one hour. After neutralizing the reaction mixture with disodium phosphate buffer (750 mM, 10 mM EDTA), the synthesized DNA was precipitated into a 5-fold excess of isopropanol to remove contaminants and to exchange the counterions to sodium. The precipitate was dissolved in MiliQ water and freeze dried. The synthesized and purified strands were stored at -20 °C until further use. The purity of the obtained oligonucleotides was confirmed with analytical HPLC.

##### *3.5 DNA annealing*

All DNA strands were used as received. All the sequences are provided in Supplementary Table S1 and Table S2. The DNA strands received from IDT and Biomers were dissolved in TE buffer (10 mM Tris-HCl, pH = 8.0) to prepare a stock solution of 1 mM and stored at -20 ˚C for further use. The complementary DNA strands intended for double stranded complexes, i.e., Complex 1 and Substrate 1 were dissolved in annealing buffer (10 mM Tris-HCl, 50 mM NaCl, 10 mM $MgCl_2$, pH = 8.0) with the same stoichiometry at -20 °C overnight to prepare a stock solution of 0.125 mM.

##### *3.6 Cell Culture*

HeLa cell line (ACC 57) and NF-kB/293/GFP-Luc Transcriptional Reporter Cell Line (TR860A-1) were purchased from Leibniz Institute DSMZ-German Collection of Microorganisms and Cell Cultures GmbH and BioCat GmbH respectively and cultured according to the guidelines. Briefly, cells were cultured in DMEM supplemented with 10% fetal bovine serum (FBS) (*v/v*) and penicillin/streptomycin in an incubator with 5% $CO_2$ for culturing and for the experimental setup, cells were adapted to DMEM supplemented with 1x insulin selenium and transferrin (ITS) and penicillin/streptomycin (Cell Buffer).

##### *3.7 Comparison of Cell Buffer and FBS-supplemented Media*

The degradation kinetics of a DNA duplex labelled with a fluorophore/quencher pair in Cell Buffer (insulin-transferrin-selenium (ITS)-containing media) and 10% FBS cell culture media (conventional media) were compared at 37 °C, by measuring the fluorescence intensity every hour. We identified that Cell Buffer is efficient in maintaining DNA stability in the cell environment for at least one day, whereas FBS-containing media shows significant degradation within 4 hours (Figure S3). This implies that the Cell Buffer provides a more stable, nuclease free environment for DNA and thus, was used for all our cell experiments.

##### *3.8 Conjugation of TNFα to DNA*

DNA was conjugated to TNF by using a previously established method[4]. 50 μM Human TNFα solution in 1x PBS was reacted with 4 equivalents of Azido-PEG12-NHS Ester for 1 hour at room temperature. Excess linker was removed using a NAP-10 desalting column.

Next, 8 equivalents of DBCO-terminated DNA (S-DBCO) were added to the TNFα-$N_3$ solution, and the reaction proceeded for 16 hours at room temperature. The TNFα-DNA conjugate was then purified and concentrated using 10K MWCO spin filter. The product was confirmed by HPLC (Figure S16).

##### *3.9. Preparation of DNA-functionalized SCs for transient release of Signal*

SC1/x-S*Q/S-Atto 647N is employed for checking FI changes by using x-S*Q instead of x-S* while keeping all conditions for annealing same as (Figure a-b). SC1/x-S*Q/S-Atto 647N is suspended in DNA-Cell buffer in a total volume of 50 μL at a final SC concentration of 0.05 wt% containing 20 μM Complex 1, 5 μM Substrate 1, 10 μM Input 1 and Input 2, 0.8 WU $\mu L^{-1}$ of T4 DNA ligase and 0.8 U $\mu L^{-1}$ of BsaI at 37 °C fueled by 40 μM ATP.

##### *3.10 ATP-fueled signal transfer from SCs to Cells*

HeLa cells were seeded overnight with a seeding density of $10^4$ cells/well in Cell Buffer. The next day cells were treated with a combination of 20 μM Complex 1, 5 μM Substrate 1, 10 μM Input 1 and Input 2, 0.8 WU $\mu L^{-1}$ of T4 DNA ligase and 0.8 U $\mu L^{-1}$ of BsaI and 0.05 wt% of SC1/x-S*/S-Atto 647N suspended in DNA-Cell buffer in a total volume of 50 μL, the system was incubated together at 37 °C and fueled by 40 μM ATP.

##### *3.11 Flow Cytometer*

HeLa cells were seeded overnight with a seeding density of $10^5$ cells/well in Cell Buffer. The next day cells were treated with a combination of 20 μM Complex 1, 5 μM Substrate 1, 10 μM Input 1 and Input 2, 0.8 WU $\mu L^{-1}$ of T4 DNA ligase and 0.8 U $\mu L^{-1}$ of BsaI and 0.05 wt% of SC1/x-S*/S-Atto 647N suspended in DNA-Cell buffer in a total volume of 50 μL, the system was incubated together at 37 °C and fueled by different ATP concentrations: 40 μM, 80 μM, 120 μM. The cells were fixed at different time points: 0.5 h, 2 h, 4 h, washed three times in PBS and resuspended in PBS before running through flow cytometer. The gating procedure is illustrated in Figure S17, and data was analyzed using FLOWJO (v10.8.1) software.

##### *3.12 ATP-fueled release of TNFα-linked ssDNA Signal*

TNFα-RCL was seeded overnight with a seeding density of $10^5$ cells/well in Cell Buffer. The next day cells were treated with a combination of 20 μM Complex 1, 5 μM Substrate 1, 10 μM Input 1 and Input 2, 0.8 WU $\mu L^{-1}$ of T4 DNA ligase and 0.8 U $\mu L^{-1}$ of BsaI and 0.05 wt% of SC1/x-S*/S-TNFα suspended in DNA-Cell buffer in a total volume of 50 μL, the system was incubated together at 37 °C and fueled by 40 μM ATP concentration. The cells were fixed 16 h after treatment, stained with CellTrace™ dye and imaged on CLSM.

#### 4. Supplementary Notes

##### *4.1 Definition of activity units of both enzymes*

**Definition of the Weiss Unit to describe the activity of T4 DNA ligase**: 0.01 Weiss Unit [WU] of T4 DNA Ligase is the amount of enzyme required to catalyze the ligation of greater than 95 % of 1 μg of λ/HindIII fragments at 16 °C in 20 minutes.

**Unit definition to describe the activity of BsaI**: One Unit [U] is defined as the amount of enzyme required to completely digest 1 μg of pXba DNA in one hour at 37 °C in 50 μl assay buffer containing acetylated BSA added to a final concentration of 0.1 g/L

#### 5. Supplementary Tables

**Table S1.** DNA sequences for the oligonucleotides synthesized following a well-defined procedure (section 3.5) with their abbreviations and modifications.

| Name | Sequence 5'→ 3' | Figure | Modification |
|---|---|---|---|
| $NH_2$-x* ($T_{20}$-x*) | TTTTTTTTTTTTTTTTTTTTTGAACCCGTATATCTATCCTA | Fig. 3-4 | 5' Amino Modifier C6 dT |

**Table S2**. DNA sequences for the oligonucleotides with their abbreviations, sequences, and modifications. 5' Phosphorylated DNA strands were purchased from IDT. All other DNA sequences were purchased from Biomers.

| Name | | Sequence 5'→ 3' | Figure | Modification |
|---|---|---|---|---|
| Substrate 1 | Output | CGGATTGGTATTGTATTACC | Fig. 2-4 | None |
| | Substrate1b | AATCTTTAATACAATACCAATCCGATT | Fig. 2-4 | 5' Phos |
| Complex 1 | Complex1a | CATGAGAATTCCATTCACGGTCTCT | Fig. 2-4 | None |
| | Complex1b | GATTAGAGACCGTGAATGGAATTCTCATG | Fig. 2-4 | 5' Phos |
| Input 1 | | AATCAATCGGA | Fig. 2-4 | 5' Phos |
| Input 2 | | GTATTAAA | Fig. 2-4 | None |
| x-S*Q (Linker) | | **TCAGGTAATACAATACCAATCCGTAGGATAGATATACGGGTTC** | Fig. 2 | 5' BMNQ 620 |
| x-S*Atto 647N (Linker) | | **TCAGGTAATACAATACCAATCCGTAGGATAGATATACGGGTTC** | Fig. S3 | 5' ATTO 647N |
| x-S* (Linker) | | **TCAGGTAATACAATACCAATCCGTAGGATAGATATACGGGTTC** | Fig. 3,4 | None |
| S-Atto 647N (Signal) | | **GGTATTGTATTACCTGA** | Fig. 4a,4b,4c | 3' ATTO 647N |
| S-Cy5 (Signal) | | **GGTATTGTATTACCTGA** | Fig. 4a,4b,4c | 3' Cy5 |
| S-Atto 488 (Signal) | | **GGTATTGTATTACCTGA** | Fig. 3,4d,4e,4f,4g | 3' ATTO 488 |
| S-DBCO | | **GGTATTGTATTACCTGA** | Fig. 5 | 3' DBCO |

### 6. Supplementary Figures

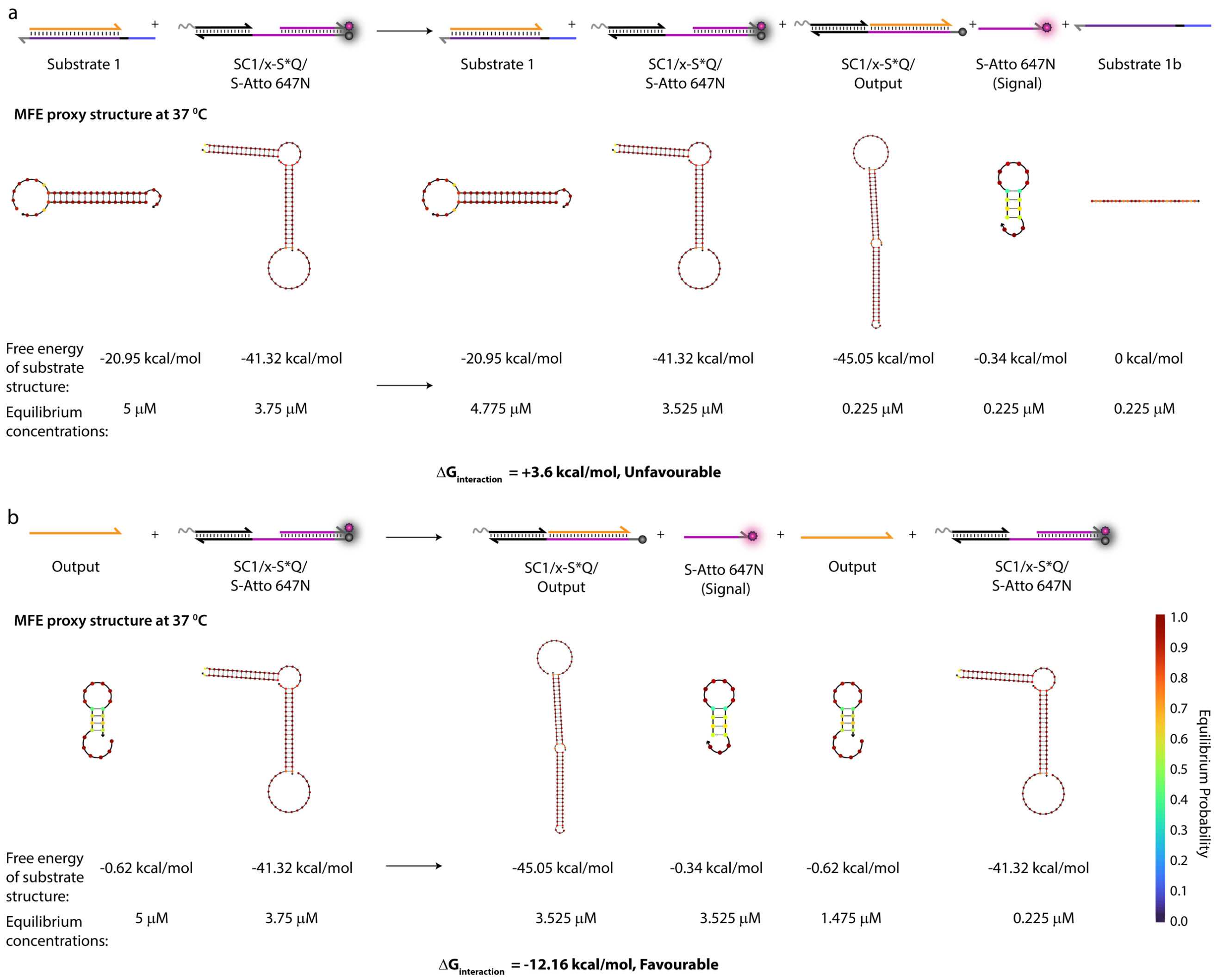

*Figure S1. Free energy change for the interaction between (a) Substrate 1 and Particle1/x-S*Q/S-Atto 647N, (b) Output and Particle1/x-S*Q/S-Atto 647N calculated with NUPACK simulations setting the temperature at 37 °C and salt concentrations at 50 mM NaCl, 10 mM $MgCl_2$. Only free Output favorably releases the Signal strand from Particle1/x-S*Q/S-Atto 647N and the interaction between Substrate and Particle1/x-S*Q/S-Atto 647N is weakly unfavorable.*

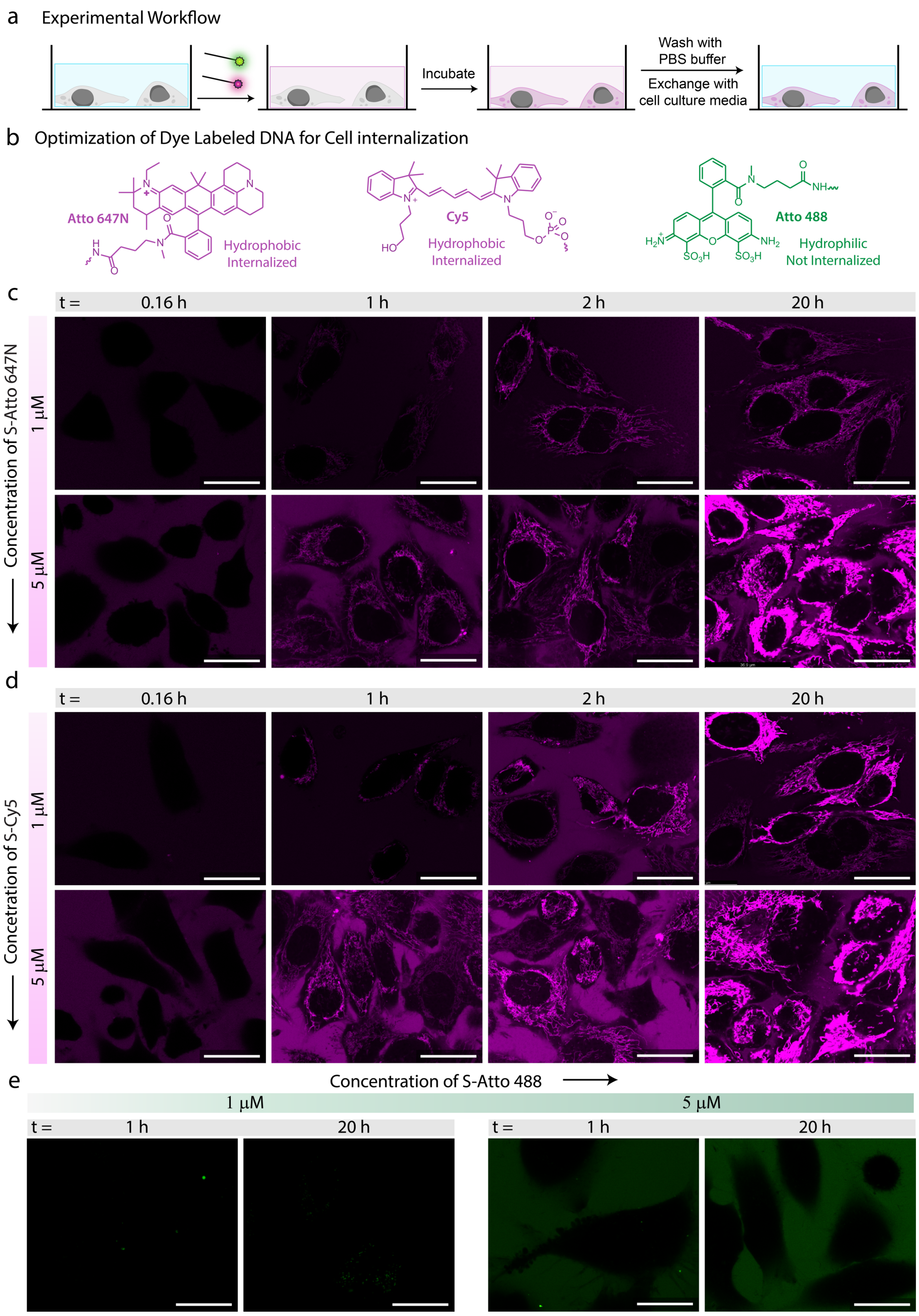

*Figure S2. Screening of different dye labelled Signal (S) strand for internalization in living cells. (a) Schematic representation of the experimental workflow where HeLa cells are incubated with respective Signal strands in DNA-Cell buffer for 24 h after which the medium is exchanged with PBS buffer. (b) Signal strands modified with Atto 647N, Cy5 and Atto 488 are used. The internalization of Signal strand depends on the hydrophobic versus hydrophilic nature of the dyes used for modification of Signal strand. (c), (d) in situ CLSM images of cells incubated with two different concentrations (1 and 5 µM) of S-Atto 647N (a) and S-Cy5 (b) at 0.16 h, 1 h, 2 h, and 20 h. (e) in situ CLSM images of cells incubated with two different concentrations (1 and 5 µM) of S-Atto 488 at 1 h and 20 h. Experimental conditions: HeLa cells at a concentration of $10^4$ cells/mL were incubated in DNA-Cell buffer at 37 °C with respective dye labelled Signal strand. Scale bars: (a), (b), and (c) 25 µm.*

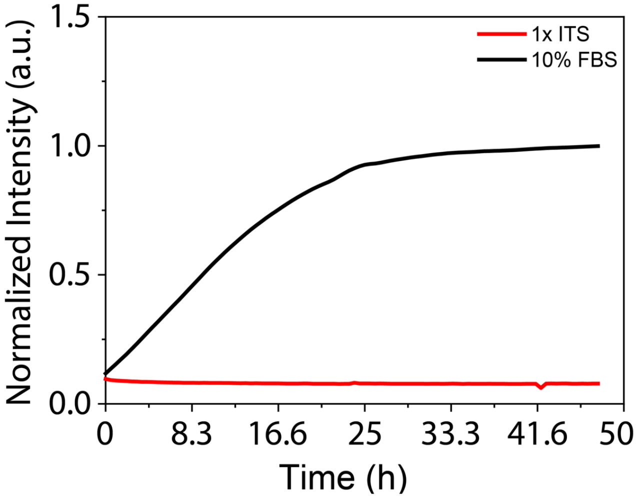

*Figure S3. Degradation kinetics of a DNA duplex labelled with a fluorophore/quencher pair in Cell Buffer (1x ITS cell culture media) and 10% FBS cell culture media (conventional media) at 37 °C. The fluorescence increases stems from degradation of the DNA duplex and separation of the fluorophore/quencher pair. The data is normalized between [0,1].*

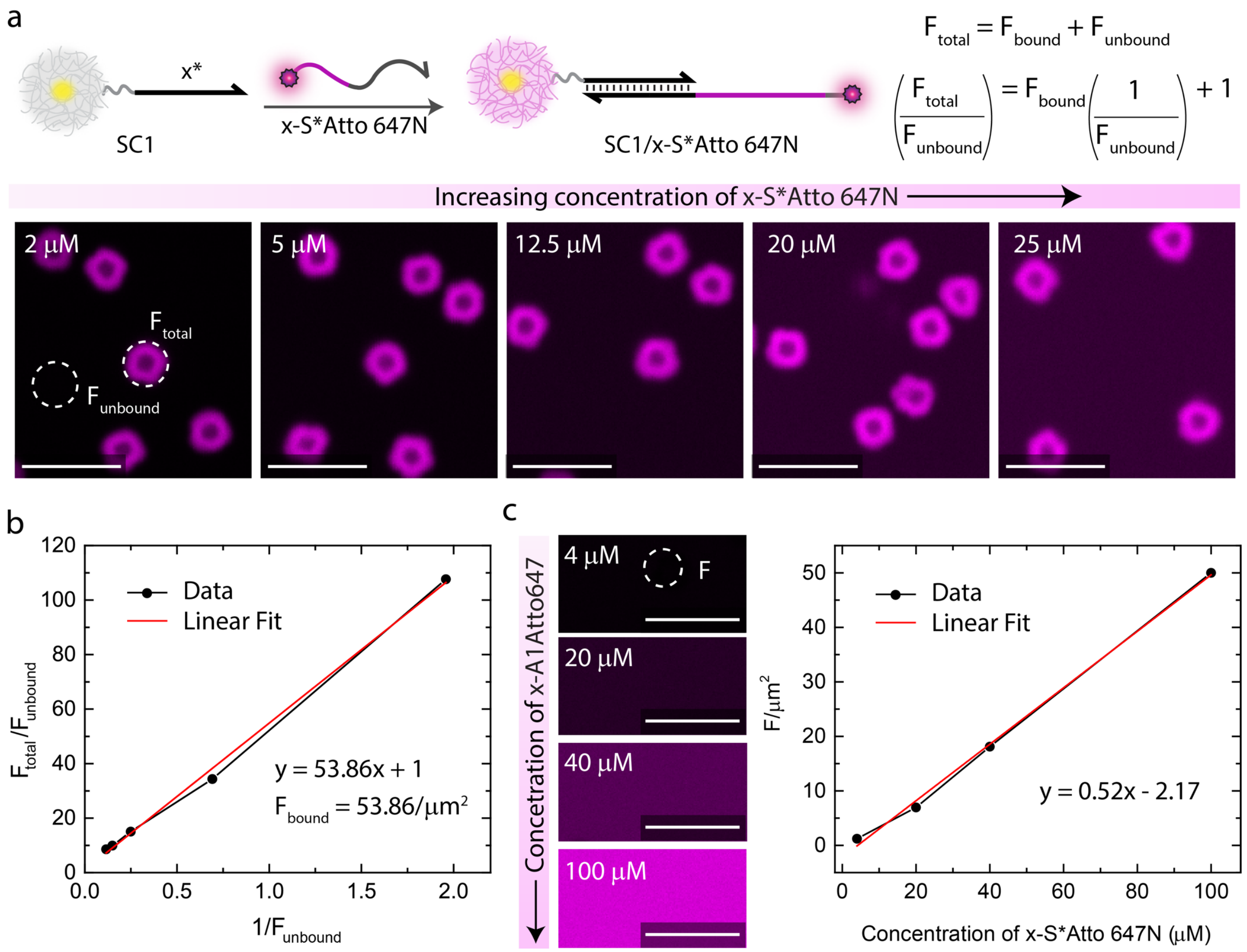

*Figure S4. Measurement of DNA grafting density on Particle1. (a) The binding capacity of x-S*Atto647 onto Particle1 via x/x* hybridization is measured as a proxy for the DNA grafting density. The mean fluorescence intensity per µm2 over the particle ($F_{total}$, includes the contribution from both x-S*Atto647 bound on Particle1 and free x-S*Atto647 in the suspension) and in the background ($F_{unbound}$, includes only free x-S*Atto647 in the suspension) is measured for increasing amounts of x-S*Atto647 via CLSM. Experimental conditions: Particle1 is incubated with increasing concentrations of x-S*Atto647 (2-25 µM) in TE buffer (pH = 8.0) at 15 °C at a final MG concentration of 0.05 wt %. $F_{total}$ and $F_{unbound}$ represent average fluorescence intensity measured from 5 different regions. (b) The data is fitted with linear equation where the slope provides fluorescence contribution from x-S*Atto647 on the particle ($F_{bound}$). (c) With the help of calibration curve between mean fluorescence intensity per µm2 (F, measured via CLSM) and concentration of free x-S*Atto647, a corresponding DNA concentration for $F_{bound}$ is calculated to be 53.86 $\pm$1.64 µM accounting for 3.3 × $10^6$ strands/MG. Experimental conditions: Increasing concentrations of z-A2-Atto647 (4-100 µM) dispersed in TE buffer. F represents an average fluorescence intensity from 5 different regions. Scale bars: (a), (c) 5 µm.*

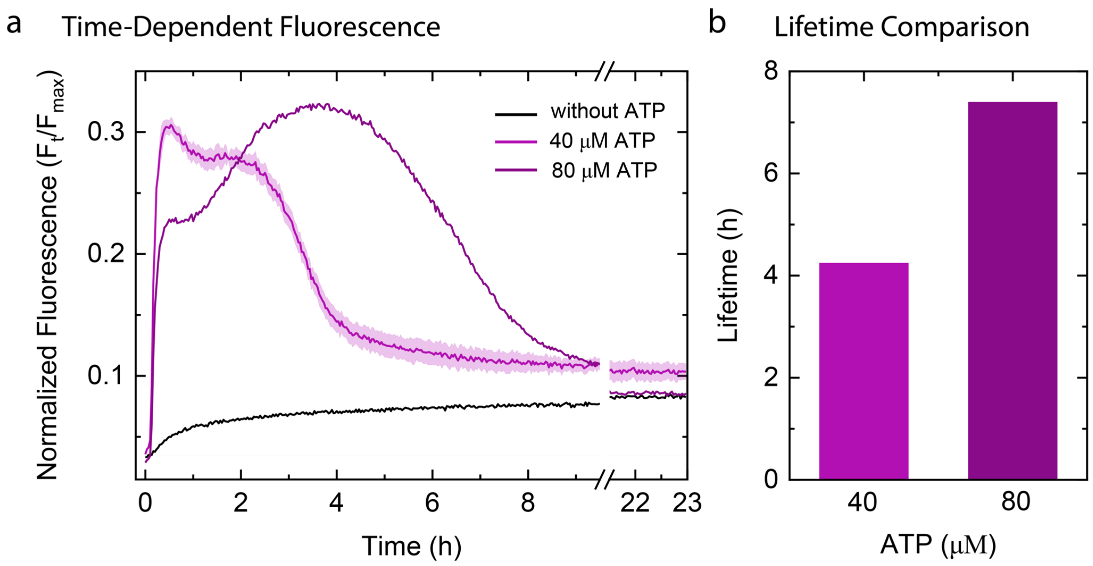

*Figure S5. ATP-dependent transient release of Signal Strand (a) Time dependent FI changes demonstrating transient increase of fluorescence because of Signal strand release into the medium upon ATP addition. The results are normalized with respect to 3.75 μM of S-Atto 647N (Signal) strands dissolved in DNA-Cell buffer which corresponds to the maximum fluorescence that can be observed in the system. The results represent an average contribution from two measurements and shaded region depict the standard deviation. (b) Corresponding lifetime obtained from (a) shows an increase with higher ATP equivalent. Experimental conditions: Particle1/x-S*Q/S-Atto 647N is employed for checking FI changes by using x-S*Q instead of x-S* while keeping all conditions for annealing same as (a), (b). Particle1/x-S*Q/S-Atto 647N is suspended in DNA-Cell buffer at a final MG concentration of 0.05 wt% containing 20 μM Complex 1, 5 μM Substrate 1, 10 μM Input 1 and Input 2, 0.8 WU μL$^{-1}$ of T4 DNA ligase and 0.8 U μL$^{-1}$ of BsaI at 37 °C fueled by different ATP concentrations (a, b).*

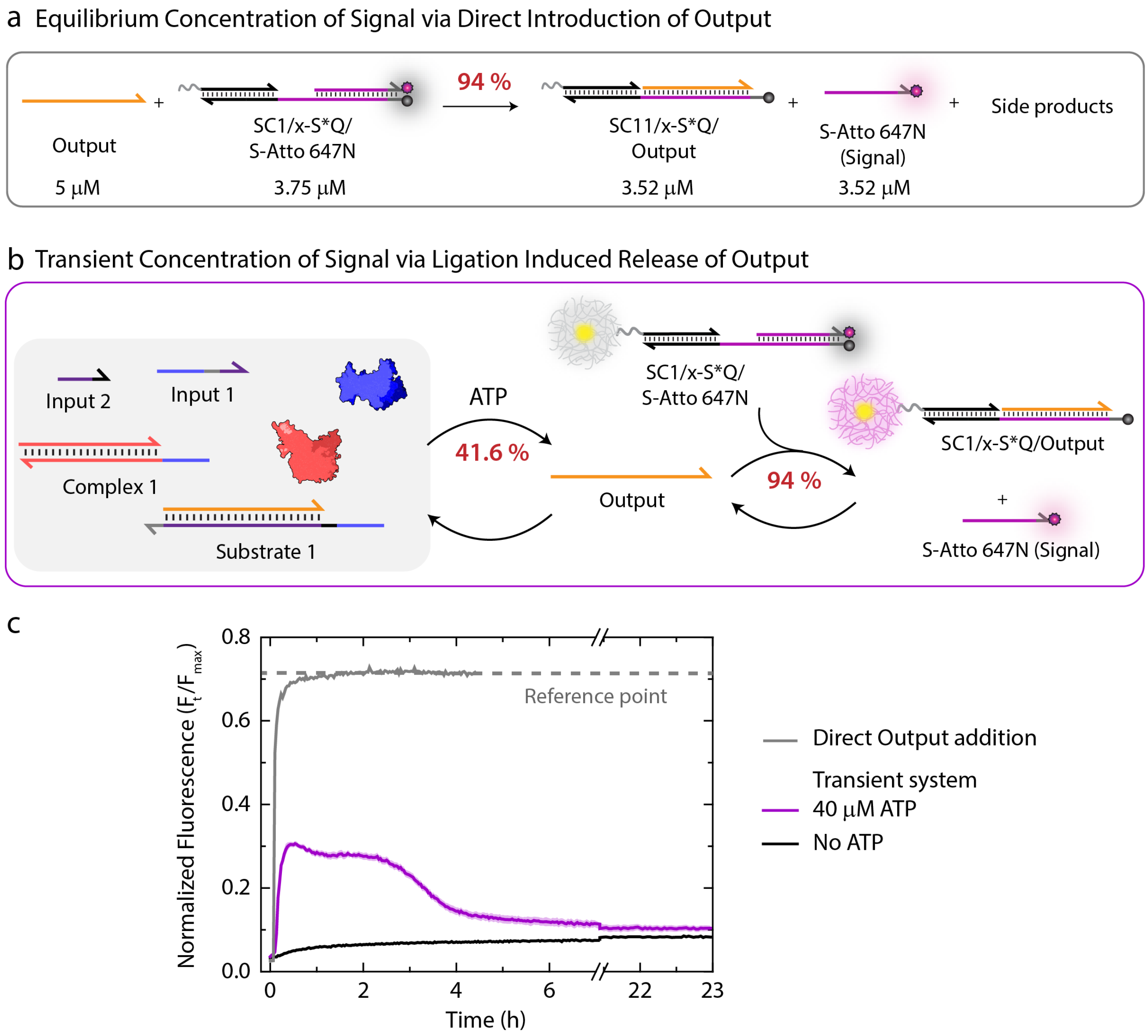

*Figure S6. (a) Equilibrium concentrations of all components obtained from NUPACK simulations upon mixing 5 µM of Output and 3.75 µM of Particle1/x-S*Q/S-Atto 647N setting the temperature at 37 °C and salt concentrations at 50 mM NaCl, 10 mM $MgCl_2$. (b) Schematics for experimental verification of percentage yield of S-Atto 647N (Signal) strand released in a transient system where Output is produced in situ via ligation. (c) Time-dependent fluorescence intensity changes demonstrating release of Signal strand when Output is added directly and when it is produced transiently via ligation. The percentage of fluorescence increase in case of direct Output addition provides a reference point which indicates the maximum fluorescence that can be observed in the transient system. This maximum fluorescence corresponds to 3.52 µM of Signal strand. In case of transient system, restriction can set in already at the hemi-ligated intermediate (e.g., Substrate 1, Complex 1 and only one of the Inputs) without completing to the fully ligated state which is the condition for expulsion of Output. Because of this, fluorescence intensity decreases by only 30% which indicates that Output is released with 41.6% efficiency with respect to Reference point. Since 94% of Output released can generate free S-Atto 647N (Signal) (a), a final yield of 39% (1.47 µM) can be attributed to free S-Atto 647N (Signal). Experimental conditions: For transient system (magenta curve), Particle1/x-S*Q/S-Atto 647N at concentration of 3.75 µM are dissolved in 1X NEB CutSmart buffer containing 20 µM Complex 1, 5 µM Substrate 1, 10 µM Input 1 and Input 2, 0.8 WU $\mu L^{-1}$ of T4 DNA ligase and 0.8 U $\mu L^{-1}$ of BsaI at 37 °C fueled by 40 µM ATP. For case with Direct Output addition (grey curve), Particle1/x-S*Q/S-Atto 647N at concentration of 3.75 µM is dissolved in 1X NEB CutSmart buffer at 37 °C followed by addition of 5 µM of Output.*

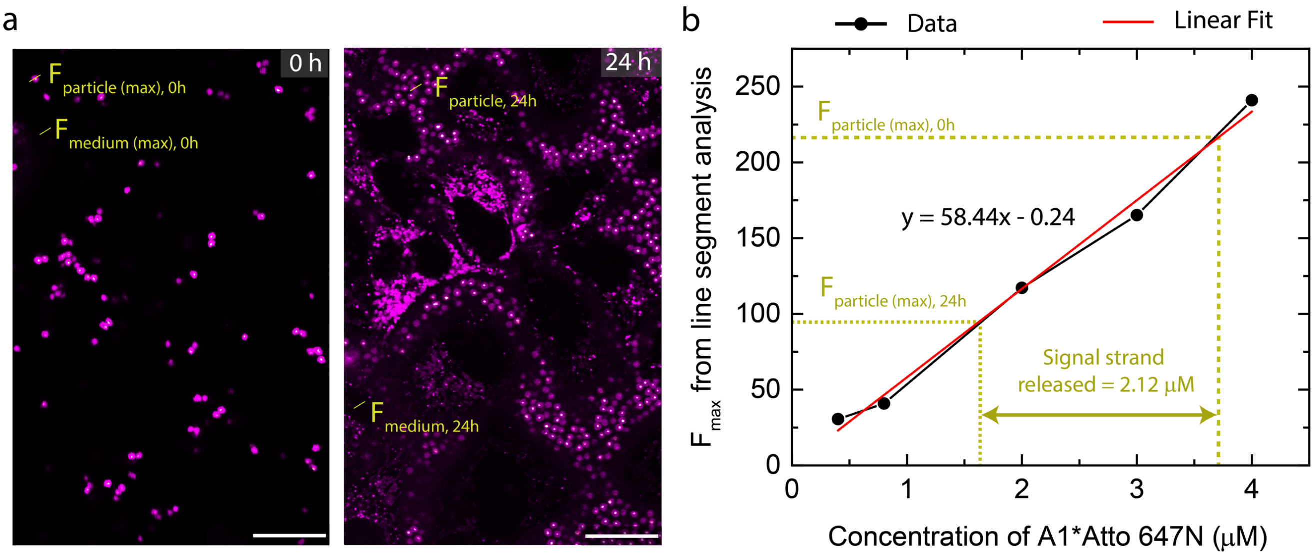

*Figure S7. Calculation of the amount of S-Atto 647N (Signal) strand released from Particle1/x-S*Q/S-Atto 647N into medium and delivered into cells after 10 h via controlled out-of-equilibrium delivery method presented in the paper (Figure 4). (a) The maximum fluorescence intensity obtained from the line segment analysis of particle at 0 h ($F_{particle(max)}$, 0h) and 10h ($F_{particle(max)}$, 10h) is correlated with the calibration curve between fluorescence intensity and concentration of free S-Atto 647N (b). The correlation indicates the concentration of Signal strand released from the particle to be 0.73 µM. Similarly, the maximum fluorescence intensity obtained from the line segment analysis of medium at 0 h ($F_{medium(max)}$, 0h) and 10h ($F_{medium(max)}$, 10 h) can be correlated with the calibration curve to acquire the concentration of Signal strand released into the cells to be 0.09 µM. Finally, the amount of Signal strand delivered into the cells comes out to be 0.64 µM or 3.85 x $10^{10}$ Signal strands per cell. Scale bars: (a) 20 µm.*

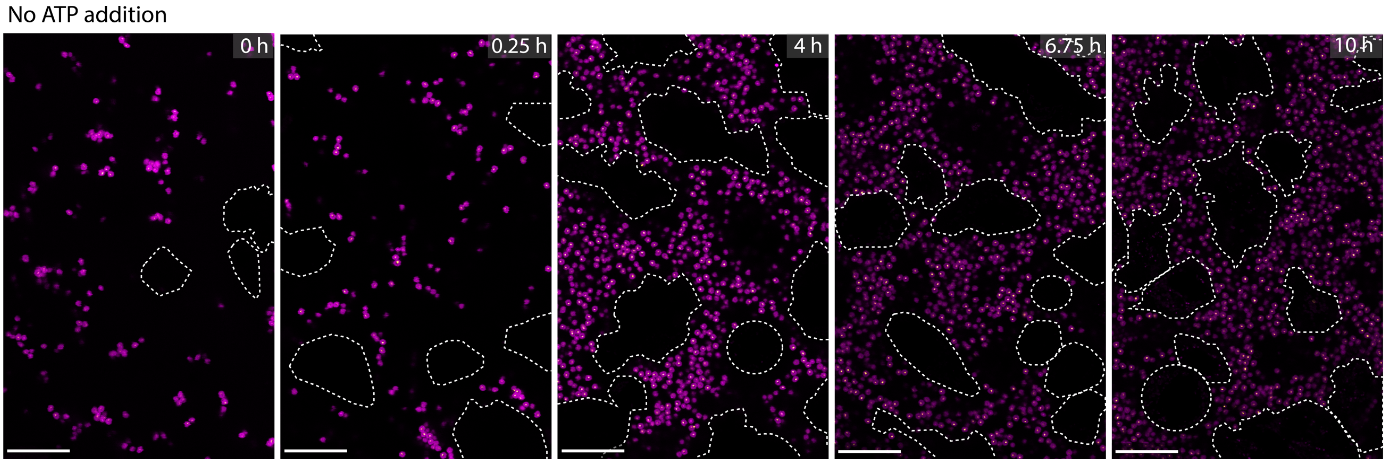

*Figure S8. CLSM imaging of the control system combining all three layers together within extracellular medium without ATP addition. Experimental conditions: Particle1/x-S*/S-Atto 647N suspended in DNA-Cell buffer at a final MG concentration of 0.05 wt% containing 20 µM Complex 1, 5 µM Substrate 1, 10 µM Input 1 and Input 2, 0.8 WU µL$^{-1}$ of T4 DNA ligase and 0.8 U µL$^{-1}$ of BsaI and HeLa cells at a concentration of 10$^4$ cells/mL incubated together at 37 °C (b, d). Scale bars: 20 µm.*

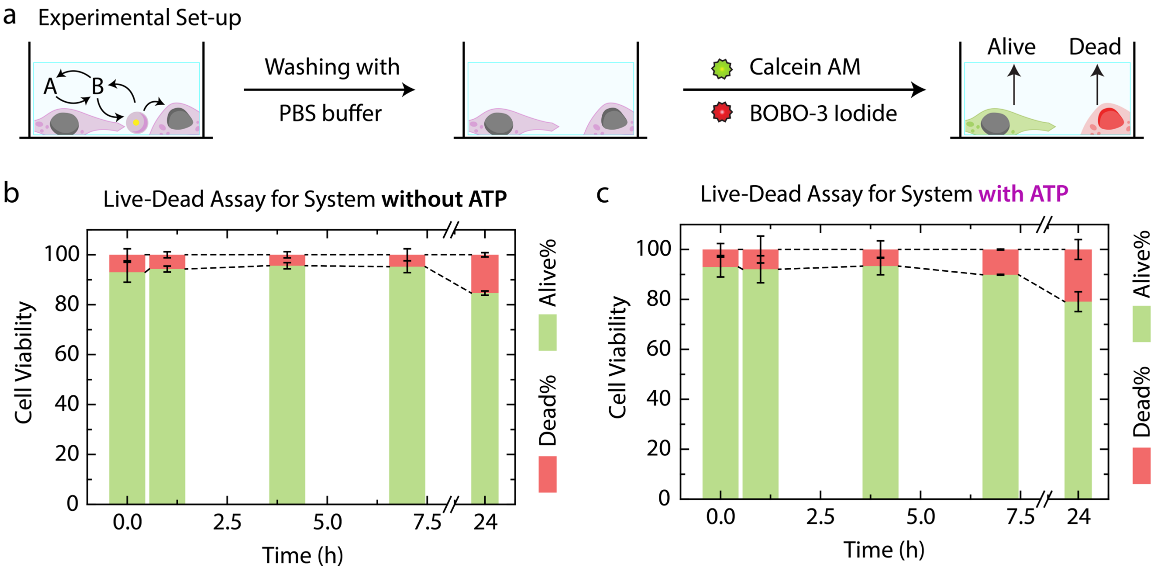

*Figure S9. Cell viability test for the transient delivery of Signal strands in DNA-Cell Buffer. (a) Schematic representation of the experimental workflow where the extracellular medium containing Layer 1 and Layer 2 is exchanged with PBS buffer followed by introduction of two different dyes, Calcein AM and BOBO-3 Iodide. Calcein AM only stains the live cells and is based on intracellular esterase activity, whereas BOBO-3 Iodide specifically penetrates the dead cells. (b), (c) Cell viability test on the samples containing all three Layers at different time points in the absence (b) and presence (c) of ATP. Five duplicate samples were prepared for five separate measurements., whenever needed, the extracellular medium was replaced with PBS buffer, equimolar concentrations of Calcein AM and BOBO-3 Iodide were added (as per the guidelines of the supplier), the sample was incubated for 15 min at 25 °C followed by acquiring microscopic images using FITC and TRITC filters for separate dyes. The images were manually analyzed to calculate the percentage of live and dead cells. The results represent an average contribution from two different images and error bars depict the SD.*

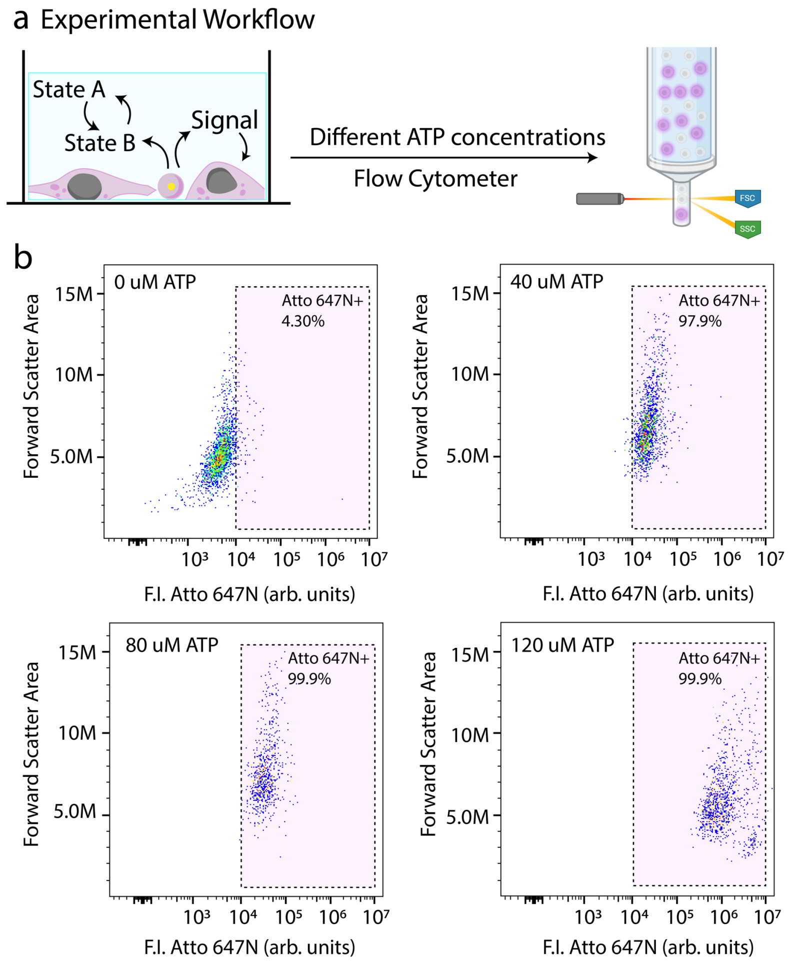

*Figure S10. Flow cytometer measurement of the cells in the presence of the ATP fueled ERN networks. (a) Schematic representation of the experimental workflow. (b) Dot plots representing the uptake of Signal strand by cells at various ATP concentrations: 0 µM, 40 µM, 80 µM, 120 µM. Experimental conditions: HeLa cells at a concentration of $10^5$ cells/well were incubated in reaction network at 37 °C, the cells were then harvested, fixed and washed with PBS at 30 min time point.*

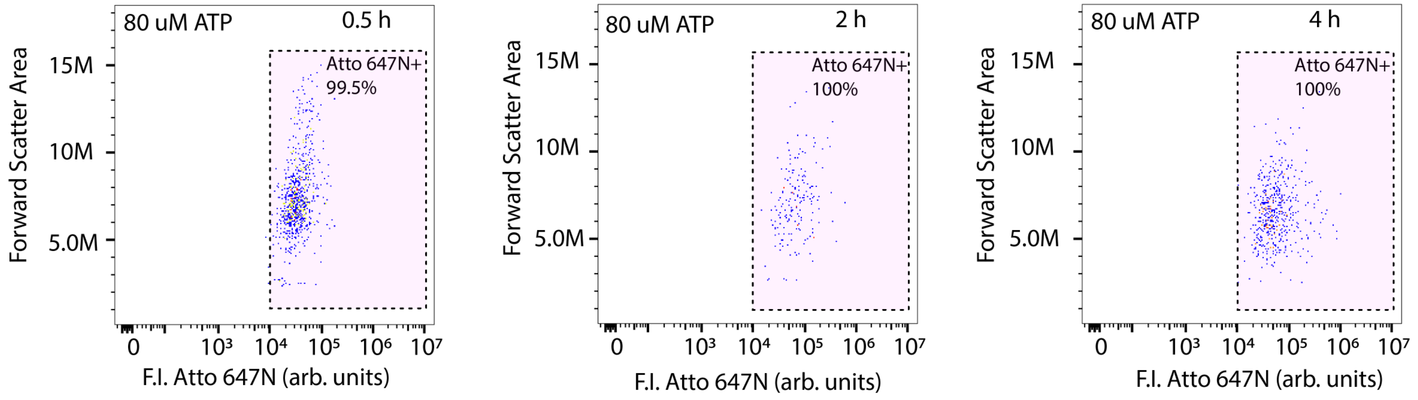

*Figure S11. Flow cytometry analysis of the cells showing dot plots with Atto 647 channel and forward scatter area on the x and y axes respectively in the presence of the ATP driven reaction network at ATP concentration of 80 µM at different time points (a) 0.5 h, (b) 2 h, (c) 4 h to study the release of dye labelled S (Signal) strand. The purple region outlined with dotted lines represents the percentage of cells that take up the Signal strand. Experimental conditions: HeLa cells at a concentration of $10^5$ cells/well were incubated in reaction network at 37 °C, the cells were then harvested, fixed and washed with PBS at 0.5 h, 2 h, 4 h time points.*

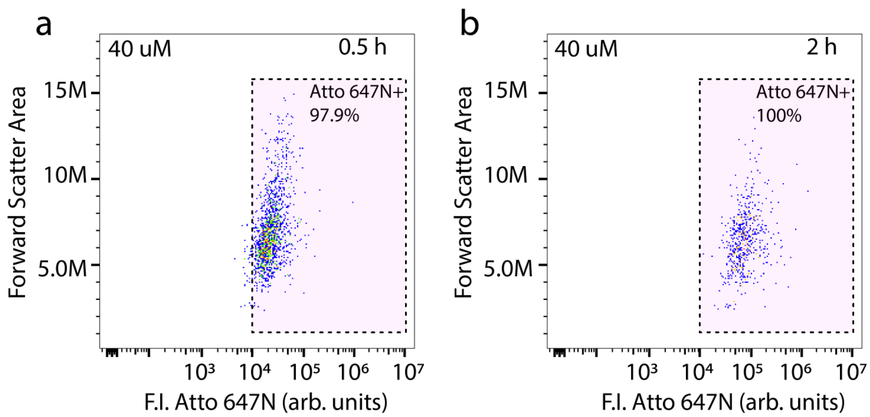

*Figure S12. Flow cytometry analysis of the cells showing dot plots with Atto 647 channel and forward scatter area on the x and y axes respectively in the presence of the ATP driven reaction network at ATP concentration of 40 µM at different time points (a) 0.5 h, (b) 2 h to study the release of dye labelled S (Signal) strand. The purple region outlined with dotted lines represents the percentage of cells that take up the Signal strand. Experimental conditions: HeLa cells at a concentration of $10^5$ cells/well were incubated in reaction network at 37 °C, the cells were then harvested, fixed and washed with PBS at 0.5 h, 2 h, 4 h time points.*

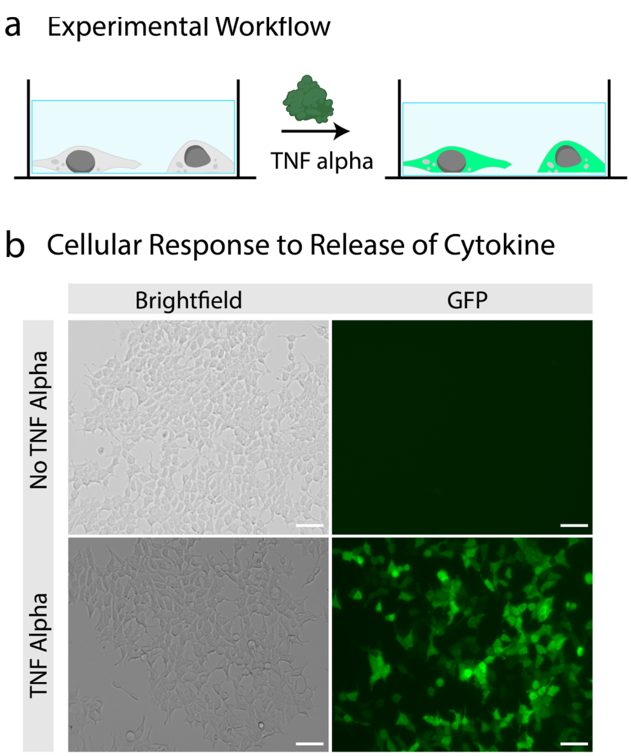

*Figure S13. Examining the effect of native TNFα on TNFα-RCL. (a) Schematic representation of the experimental workflow where native TNFα is incubated with cells. (b) Brightfield and fluorescence images of TNFα-RCL with and without TNFα incubation in DNA-Cell Buffer. Scale bar: 20 µm. Experimental conditions: TNFα-RCL seeded overnight at concentration of $10^5$ cells/ well at 37 °C, subsequently treated with 1 µM TNFα for 16 h. Imaging was performed at 16 h time point.*

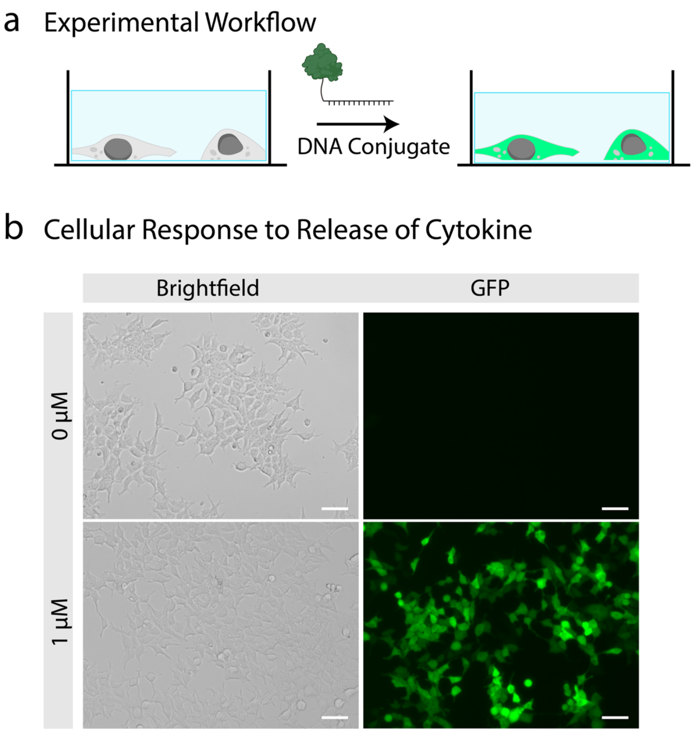

*Figure S14. Examining the effect of TNFα-DNA conjugate on TNFα*-RCL. *(a) Schematic representation of the experimental workflow where TNFα-DNA conjugate is incubated with cells. The concentration of TNFα-DNA conjugate is 0 µM, 1 µM. (b) Brightfield and fluorescence images of TNFα*-RCL incubated *with TNFα-DNA conjugate in DNA-Cell Buffer. Scale bar: 20 µm. Experimental conditions: TNFα-RCL seeded overnight at concentration of $10^5$ cells/ well at 37 °C, subsequently incubated with 0 µM and 1 µM TNFα-DNA conjugate for 16 h. Imaging was performed at 16 h time point.*

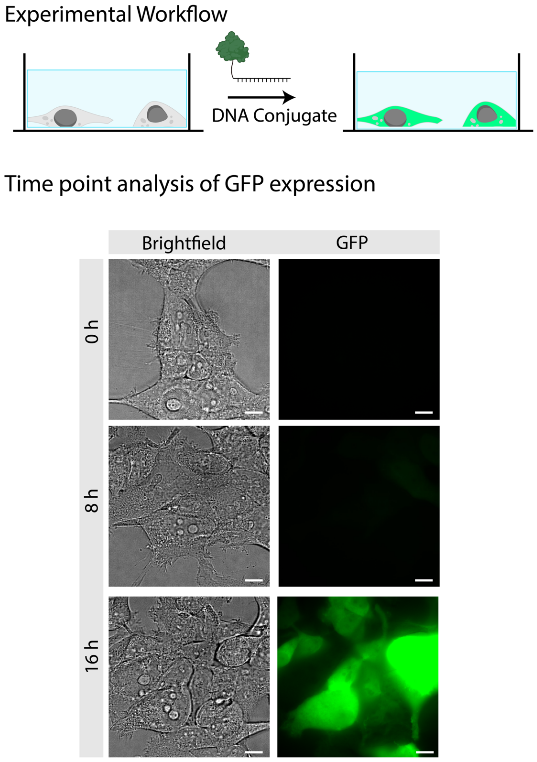

*Figure S15. Time-dependent study of GFP expression in TNFα-RCL incubated with a DNA–TNF-α conjugate in DNA-Cell Buffer. Brightfield and fluorescence images of TNFα-RCL captured at 0 h, 8 h, 16 h post treatment. Scale bar: 10 µm. Experimental conditions: TNFα-RCL seeded and incubated overnight at concentration of $10^5$ cells/ well at 37 °C, followed by treatment with 1 µM TNFα-DNA conjugate. Imaging was performed hourly following treatment.*

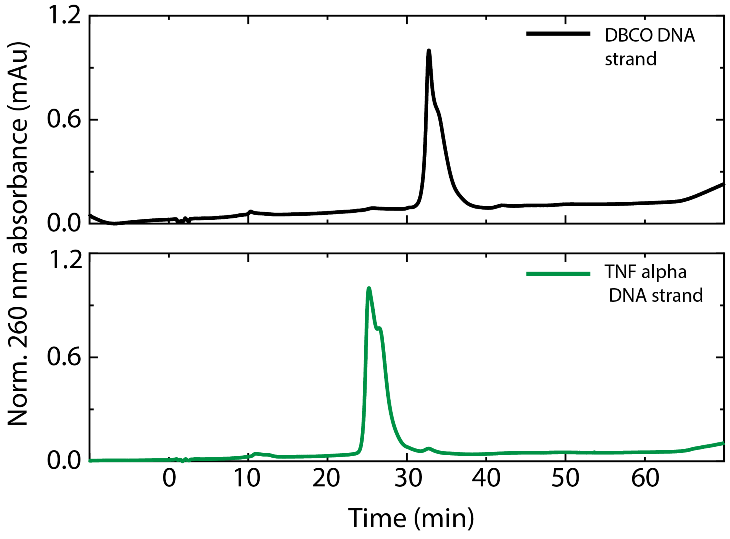

*Figure S16. Characterization of DNA conjugated with TNFα protein. HPLC graphs depicting different elution times of DNA-DBCO ssDNA and TNFα DNA ssDNA. The data is normalized between [0,1].*

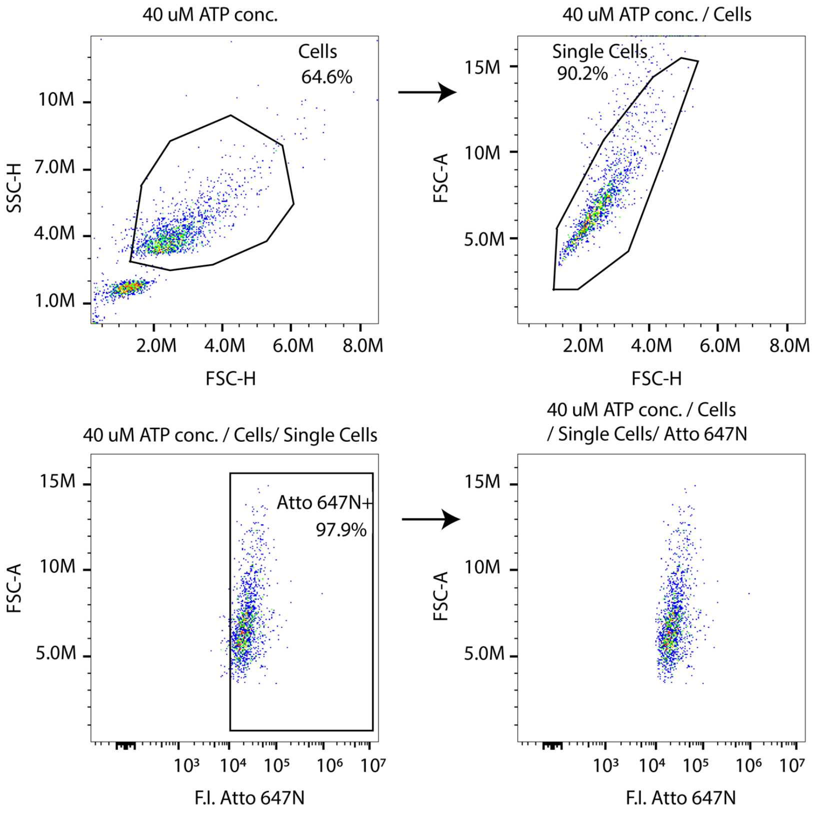

*Figure S17. Flow cytometer gating workflow illustrated through representative dot plots.*

#### 7. Supplementary References

[1] K. Han, D. Go, D. Hoenders, A. J. C. Kuehne, A. Walther, *ACS Macro Lett.* **2017**, *6*, 310–314.
[2] C. Sharma, A. Samanta, R. S. Schmidt, A. Walther, *J. Am. Chem. Soc.* **2023**, *145*, 17819–17830.
[3] Z. Meng, M. H. Smith, L. A. Lyon, *Colloid Polym. Sci.* **2009**, *287*, 277–285.
[4] N. Ramani, C. A. Figg, A. J. Anderson, P. H. Winegar, E. Oh, S. B. Ebrahimi, D. Samanta, C. A. Mirkin, *Adv. Mater.* **2023**, *35*, 2301086.
